# Pathway Commons: 2019 Update

**DOI:** 10.1101/788521

**Authors:** Igor Rodchenkov, Ozgun Babur, Augustin Luna, Bulent Arman Aksoy, Jeffrey V. Wong, Dylan Fong, Max Franz, Metin Can Siper, Manfred Cheung, Michael Wrana, Harsh Mistry, Logan Mosier, Jonah Dlin, Qizhi Wen, Caitlin O’Callaghan, Wanxin Li, Geoffrey Elder, Peter T. Smith, Christian Dallago, Ethan Cerami, Benjamin Gross, Ugur Dogrusoz, Emek Demir, Gary D. Bader, Chris Sander

**Affiliations:** The Donnelly Centre, University of Toronto, Toronto, Ontario, M5S 3E1, Canada; Department of Molecular and Medical Genetics, School of Medicine, Oregon Health & Science University, Portland, OR, 97239, USA; cBio Center, Department of Data Sciences, Dana-Farber Cancer Institute, Boston, MA, 02215, USA; Department of Cell Biology, Harvard Medical School, Boston, MA, 02215, USA; Computational Biology Center, Memorial Sloan Kettering Cancer Center, New York, NY, 10065, USA; Tri-Institutional Training Program in Computational Biology and Medicine, New York, NY, 10065, USA; Department of Systems Biology, Harvard Medical School, Boston, MA, 02215, USA; Department of Informatics, Technische Universität München, 85748 Garching, Germany; Human Oncology and Pathogenesis Program, Memorial Sloan Kettering Cancer Center, New York, New York 10065, USA; Marie-Josée and Henry R. Kravis Center for Molecular Oncology, Memorial Sloan Kettering Cancer Center, New York, New York 10065, USA; Department of Computer Engineering, Bilkent University, Ankara, 06800, Turkey

## Abstract

Pathway Commons (https://www.pathwaycommons.org) is an integrated resource of publicly available information about biological pathways including biochemical reactions, assembly of biomolecular complexes, transport and catalysis events and physical interactions involving proteins, DNA, RNA, and small molecules (e.g., metabolites and drug compounds). Data is collected from multiple providers in standard formats, including the Biological Pathway Exchange (BioPAX) language and the Proteomics Standards Initiative Molecular Interactions format, and then integrated. Pathway Commons provides biologists with (1) tools to search this comprehensive resource, (2) a download site offering integrated bulk sets of pathway data (e.g., tables of interactions and gene sets), (3) reusable software libraries for working with pathway information in several programming languages (Java, R, Python, and Javascript), and (4) a web service for programmatically querying the entire dataset. Visualization of pathways is supported using the Systems Biological Graphical Notation (SBGN). Pathway Commons currently contains data from 22 databases with 4,794 detailed human biochemical processes (i.e., pathways) and ∼2.3 million interactions. To enhance the usability of this large resource for end-users, we develop and maintain interactive web applications and training materials that enable pathway exploration and advanced analysis.

## INTRODUCTION

Pathway information that describes interactions between molecules in biological processes can help in solving research problems, such as the interpretation of genomics data (1), generating hypotheses surrounding disease mechanisms (2, 3), design of rational therapeutics (4) and treatment decision strategies (5). In a recent translational study, using pathway resources, genome-wide observations of increased methylation in pediatric brain cancer were linked to an upstream methyltransferase, which could be targeted pharmacologically and has served as the basis of clinical trials (6).

The number of available pathway and interaction resources has nearly tripled over the last decade, from 190 in 2006 to 702 in 2018 (7)(www.pathguide.org), increasing the need for integration. Unfortunately, making this knowledge available to the research community has been hindered by fragmentation from the use of diverse data representation schemes and softwares, making pathway information from multiple sources difficult to combine and use.

Pathway Commons (PC) is a resource that aggregates data from publicly available biological pathway and molecular interaction databases and provides it from a single access point on the web (8). In this way, PC facilitates integration and exchange of molecular-level descriptions of metabolic and signaling pathways, molecular and genetic interactions and gene regulation networks. Data is collected from providers in the Biological Pathway Exchange (BioPAX) Level 3 (9) and the Proteomics Standards Initiative Molecular Interaction (PSI-MI) formats (10), and stored uniformly in BioPAX format. Use of the BioPAX ontology and format enables PC to capture, in a uniform and consistent way, details concerning genes, macromolecules (proteins) and small molecules and their involvement in different types of physical interactions, such as biochemical reactions, catalysis, post-translational protein modifications, complex assembly, and transport. PSI-MI data captures molecular interactions from small and large scale experiments. These descriptions are richly annotated with links to citations, experimental evidence, and external database information, for instance, protein sequence annotation. PC aims to add value to curated source databases by normalizing, integrating and exporting data in ways that simplify usage.

PC has been used to analyze transcriptomics, proteomics, and metabolomics data in a large number of projects across diseases to further our understanding of human biology in health and disease (4, 11–18). Since our original report in 2011, significant advances have been made with regard to the breadth and volume of data available along with software tools to support data creation, validation, and accessibility in the wider research community. Here, we summarize developments made since our original report and discuss future efforts to enhance accessibility and provide scalable systems for knowledge capture in support of biomedical discovery.

## PATHWAY AND INTERACTION DATA COVERAGE

PC currently integrates data from 22 public databases, up from the 9 in our initial report. This has more than tripled the number of pathways (from 1,477 to 4,794) and interactions (from 687,883 to over 2.3 million) (Figure 1). The new data covers 18,490 genes with associated HUGO Gene Nomenclature Committee (HGNC) identifiers and 11,437 small molecules associated with records from Chemical Entities of Biological Interest (ChEBI), Human Metabolome Database (HMDB), Kyoto Encyclopedia of Genes and Genomes (KEGG) Compound, and/or DrugBank (19–22). PC focuses on collecting human pathway data since many data providers focus specifically on interactions occurring in human cells.

**Figure 1.**
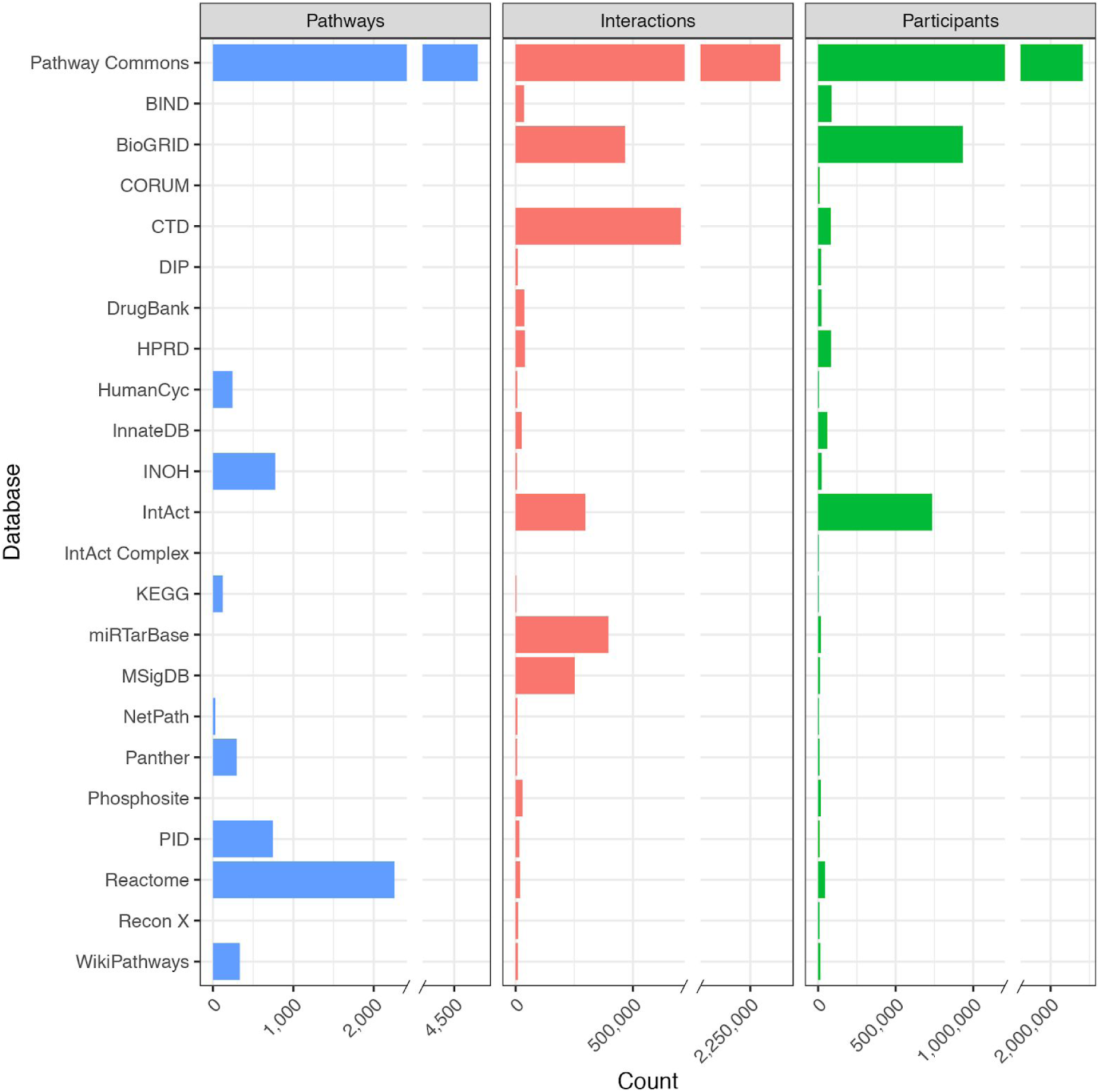
A summary of pathway and interaction databases in Pathway Commons Version 11, released February 2019. Participants are counts of “PhysicalEntity” class instances from the BioPAX ontology, which includes the classes: complexes, DNA, DNARegion, Protein, RNA, RNARegion, and SmallMolecule, including the possibility of multiple molecular states per gene (e.g. phosphorylated proteins, proteins in the nucleus). Citations for data providers: (21, 31, 62, 68–85).

## SOFTWARE INFRASTRUCTURE

The core software tools driving PC are cPath2 and Paxtools. cPath2 is an open-source database and web application for collecting, storing, and querying biological pathway data, and has been completely rewritten based on cPath (23). cPath2 is built atop the Java Paxtools library (24) which provides an in-memory BioPAX object model designed to provide an API along with rich and fast data querying, validation and format conversion utilities (25–27) (Figure 2). cPath2 includes built-in identifier mapping for linking between identical interactors and to external resources as well as an application programming interface (API) that functions as a web service for searching and retrieving pathway data sets. The web service is implemented using a RESTful architecture and allows fine-grained data retrieval as JSON-LD (to support easy access from web applications), BioPAX and other formats (see DATA FORMATS AND AVAILABILITY). It supports search, including keyword-based, in addition to the advanced querying facilities made available by Paxtools (e.g. graph-based querying). For a detailed graphical representation of pathways, Pathway Commons uses the standard Systems Biological Graphical Notation (SBGN) (28), used to reduce the ambiguity in representations of biological maps, and its accompanying SBGN-ML format (29). The web service is a major access point for software developers and computational biologists to programmatically access PC data and can be used to build third-party software apps, such as the ones described below.

**Figure 2.**
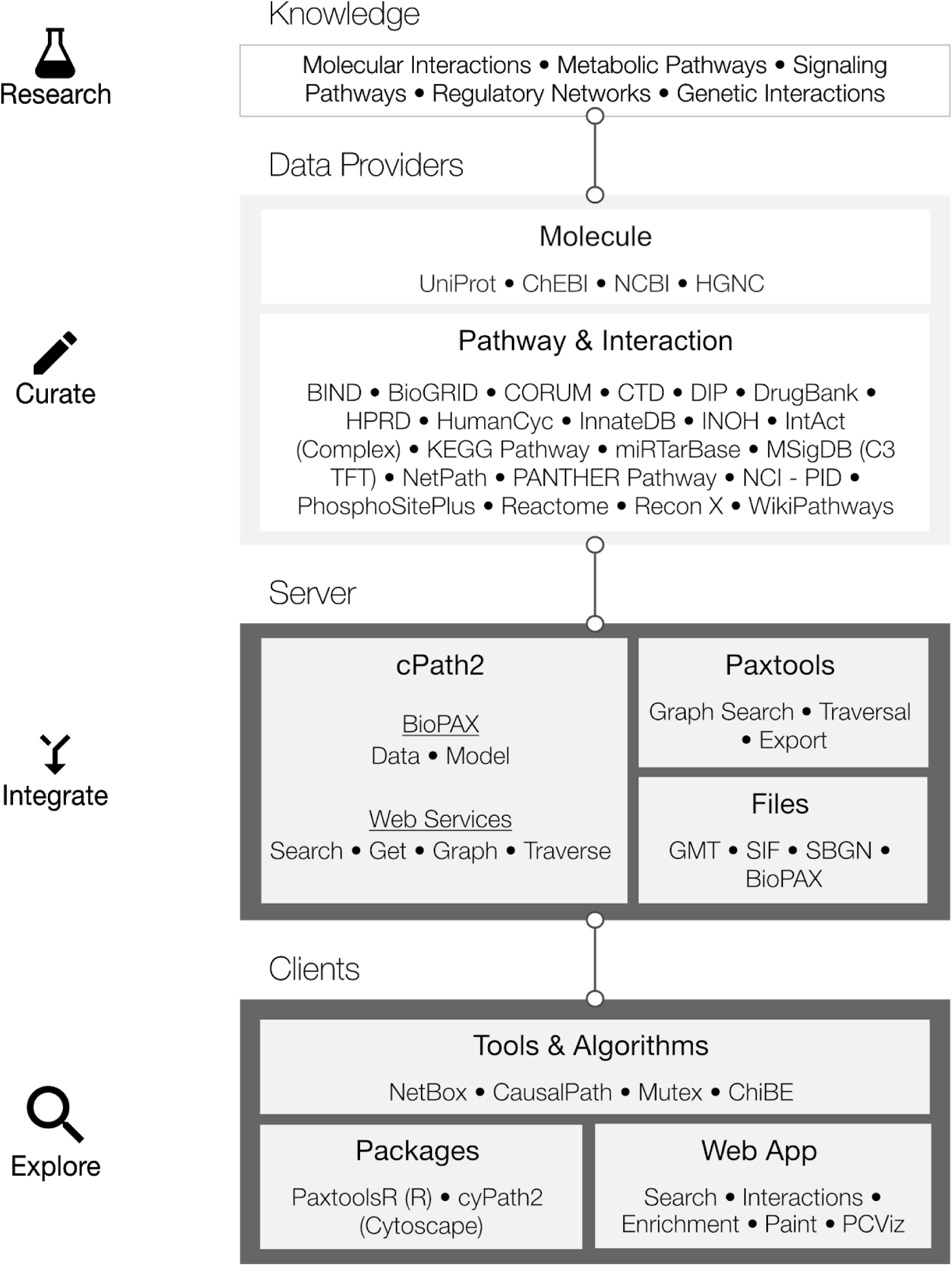
From primary knowledge to end-user pathway tools. Pathway Commons (PC) aggregates and disseminates pathway and interaction knowledge from 22 databases (version 11). BioPAX files are downloaded directly from data providers and are subsequently validated, normalized and merged into PC. Data can be directly accessed programmatically via the web service or downloaded in bulk files. Exploration and analysis are aided by software tools, packages and web apps that are tailored to the use cases of computational and experimental researchers.

## DATA FORMATS AND AVAILABILITY

Users can freely access PC data by either downloading data files (designed for computational biologists), through a web service (for software developers or computational biologists), or via a series of interactive web-based search tools. Pathway information downloads are made available in BioPAX format, Gene Matrix Transposed (GMT) format, which is used in gene set enrichment analyses (30, 31), Simple Interaction Format (SIF) and extended SIF with additional fields, which are useful for network analysis and visualization (pathwaycommons.org/pc2/formats; SUPPLEMENTARY DATA). GMT datasets are provided with HGNC or UniProt identifiers (32, 33). Users can access a file containing the entire collection or files that only contain data provided from an individual database. Data updates are scheduled approximately biannually (current release as of February 2019 is Version 11) and previous versions are also available in an archive (pathwaycommons.org/archives).

## SOFTWARE TOOLS

We have developed a number of tools using the core cPath2 and Paxtools PC infrastructure, including programming libraries as well as desktop and web-based applications for use by a broad audience.

### Tools for querying and visualizing Pathway Commons data using BioPAX

In addition to the core Java-based Paxtools library, programming libraries in other languages commonly used by computational biologists, including R (34) and Python (35), have been developed by the PC team and the community. These packages enable users to access content in BioPAX and act as clients for the PC web service. ChiBE is a desktop application focused on network visualization of BioPAX data and the analysis of genomic data in a pathway context (25, 36). Cytoscape (37) and CellDesigner (38), two widely used independent desktop tools for modeling, visualization and analysis of biological networks and pathways, have BioPAX and PC support through plugins. For instance, the Cytoscape CyPath2 plugin enables direct querying of PC from Cytoscape (apps.cytoscape.org/apps/cypath2), and the CellDesigner BioPAX export plugin allows export from Cytoscape in BioPAX format (39).

### Tools for visualizing and interacting with pathway diagrams online

A number of reusable tools have been built to enable users to interact with pathway figures online and to map data onto pathway diagrams (20, 40). We have developed software to visualize and interact with network diagrams using the SBGN standard (28). Specifically, our sbgnml-to-cytoscape and cytoscape-sbgn-stylesheet JavaScript packages (github.com/PathwayCommons) allow developers to load and style SBGN diagrams represented in the SBGN-ML plain-text format as interactive diagrams in Cytoscape.js (41). From there, figures can be exported as static images or included as part of a dynamic web application.

By virtue of exporting all pathways to SBGN, PC is able to provide a consistent visualization across all data, regardless of whether it was offered by the provider. A useful feature of PC network visualizations is automated layouts. Both SBGN exported by Paxtools and diagrams visualized in Cytoscape.js are laid out using the Compound Spring Embedder (CoSE) and fCoSE graph layout algorithms that are capable of laying out SBGN-styled pathways and complexes (graphs including nesting); the CoSE algorithm has been implemented both in Java and Javascript (26).

Together, these libraries provide the fundamental components to build rich applications to visualize pathways stored in PC and elsewhere. Examples of mature applications using these components include Newt (newteditor.org), which is a fully-featured SBGN editor that can load data from PC and other sources.

### Analytical tools using the Pathway Commons data source

A number of analysis packages that make use of PC data have been developed (42–52). Here we briefly describe several tools developed by the PC team. NetBox is an algorithm that automates the data-driven definition of network modules on the basis of genomic or molecular alterations (53). CausalPath identifies potentially causal relations between (phospho)proteomic measurements based on known pathways (54, 55). The Mutex method analyzes cancer gene alterations to detect mutual exclusivity in groups of genes, which nominates them as potential cancer drivers. Such mutual exclusivity may occur when several genes have the same downstream effect when they are altered, and altering one gene is enough for that downstream effect. Mutex uses signaling relations in PC to reduce its search space to the gene groups with a common downstream target (56). A derivative of Mutex detected which pathways are targeted by functional mutations in autism spectrum disorder *de novo* mutations (57). The Enrichment Map pathway enrichment analysis workflow incorporates all PC pathways represented as gene sets (30). PC data has also been used as prior information to predict cellular response based on data collected in systematic perturbation experiments (58). Several tools and algorithms developed within DARPA’s Big Mechanism program extensively use PC to evaluate fragments extracted from the literature using machine reading (35, 59, 60).

## WEB APPLICATIONS AND TRAINING MATERIALS

Pathway Commons maintains a number of web applications and training materials aimed at advancing pathway analysis in the research community.

### PC web apps: search and visualization

The PC search app attempts to anticipate the context of user questions from their queries and returns relevant results (apps.pathwaycommons.org/search). The system recognizes specific search types (e.g. genes) that are typically part of user queries (e.g. “*cell cycle arrest involving TP53 and CDKN1A*”). In this case, the search results display additional information about each gene along with links to additional apps that use this gene-based information as input (below) (Figure 3A). A list of pathway search hits is displayed including information about the data source and its number of “participants”. Pathway search hits link to an interactive viewer that displays the network, rendered using SBGN (see Data representations section) (Figure 3B). Clicking on any node in the visualization reveals a tooltip that contains more detailed information including type (e.g. “protein” or “Biochemical Reaction”), alternative names, supporting publications and links to other databases.

**Figure 3.**
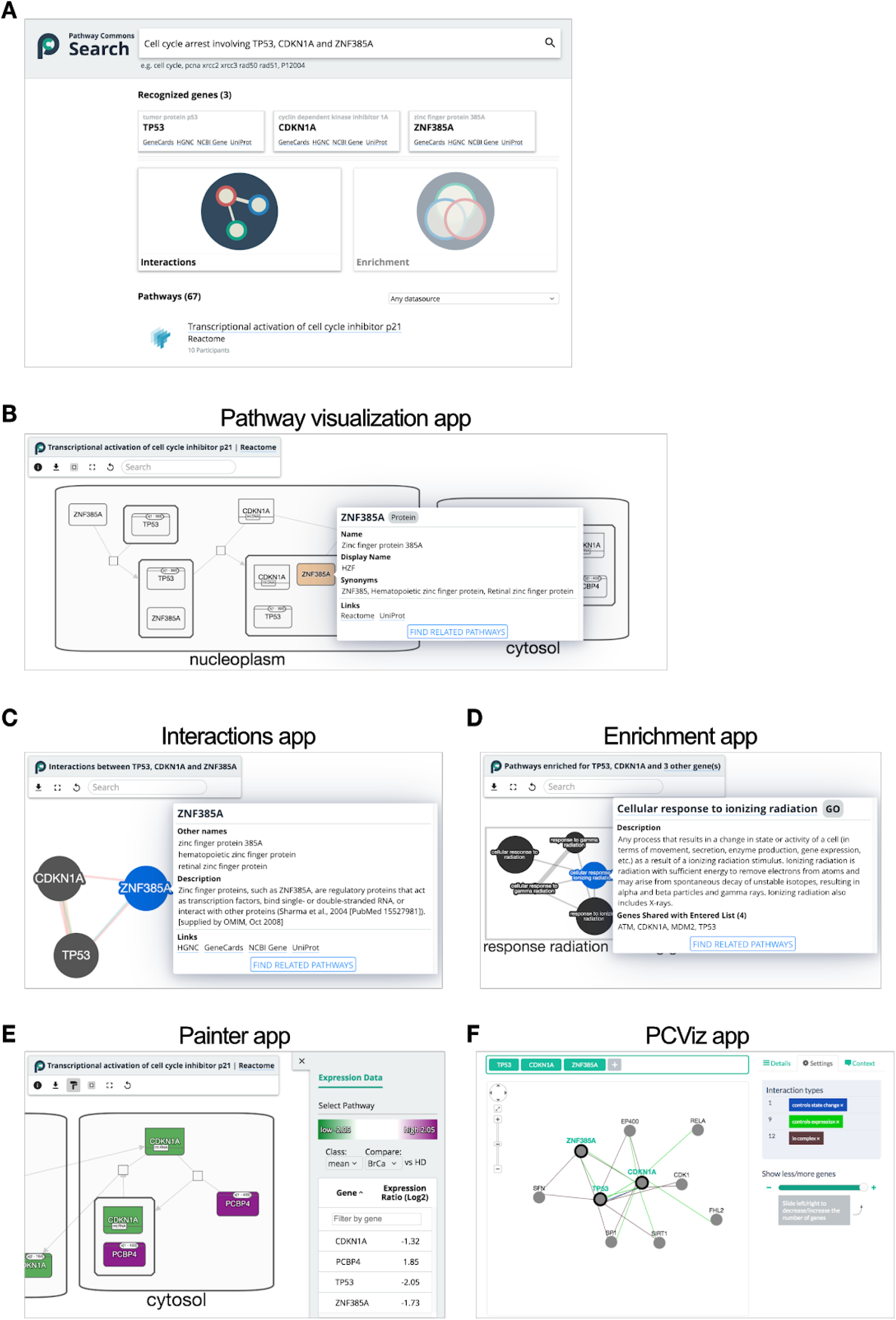
Pathway Commons web apps. **A**. Search provides integrated access to the entire collection of pathways and interactions in Pathway Commons. User queries are analyzed to select the type of search results they may find most useful, such as mentions of recognized genes along with link-outs to web apps and a ranked list of pathway search hits. **B**. Each pathway search hit is linked to an interactive viewer, rendered using the Systems Biology Graphical Notation (SBGN) visual language. **C**. The Interactions web app accessed from the search page links to an interactive network visualization showing relationships between one or more of genes recognized in a user query. **D**. With longer lists of recognized genes, an Enrichment web app links to results of pathway enrichment analysis displayed as an interactive Enrichment Map network. Nodes represent pathways (GO: Biological Process, Reactome pathways) and edges connect similar pathways, as measured by the number of shared genes. All visualization features are built using the Cytoscape.js software library. **E**. The Painter app, launched via the Enrichment Map app for Cytoscape desktop (not shown), projects quantitative gene expression data onto pathways. **F**. A PCViz app accepts one or more query genes and displays a network of interactions between and around it.

Depending on the nature of the input query, links to other apps will become available. For instance, if one or more genes are recognized in a search query, they are used to seed an interactive network visualization, called the Interactions app (Figure 3C). When the query contains one recognized gene, this app displays an interaction “neighborhood” to answer the question “What interacts with my gene?”; when multiple genes are present, only direct interactions between those genes will be shown, answering the question “How are these genes connected?”. These results are retrieved by performing either cPath2 neighborhood (one gene) or paths-between (multiple genes) web service queries. Users can filter individual interactions for specific interaction types. When the system recognizes many gene mentions, a link to the Enrichment app is enabled (Figure 3D). This app answers the question “In which pathways are the genes significantly enriched?”. Enriched pathways, computed by g:Profiler (61), are drawn from Reactome (62) and biological processes from the Gene Ontology (GO) (63). Results are displayed as a network where the nodes represent pathways containing query genes, following the enrichment map visualization concept (64). In this map the number of genes in each pathway is indicated by node size and the extent of shared genes between two pathways is indicated by the thickness of their coincident edge. To provide a high-level overview of pathways, highly overlapping pathways are clustered and labeled with terms frequently found in their pathway names. The Painter app enables annotating a pathway with gene expression data, coloring each gene according to its expression score (Figure 3E). The Painter app can be opened from an Enrichment Map result in the Cytoscape desktop app.

An additional web-based network visualization app called PCViz, helps users obtain details about genes and their interactions from PC. When queried with one or more gene or protein identifiers, PCViz displays an interaction neighborhood both between and surrounding the query genes (Figure 3F). Interactions are filterable by type and gene-gene co-citations. For biological entity nodes, a brief description and links to other biological databases are available. For interactions, the primary data source and links to publications are listed. A “context” tool enables users to display networks relevant to cancer studies by loading in data from cBioPortal (65). Downloads of the resulting network are available in PNG, SIF, and BioPAX formats.

All network views in the above-described web apps are implemented using the Cytoscape.js graph visualization JavaScript library (41).

### Training

A major goal of Pathway Commons is to support the analysis and interpretation of molecular and genomic profiling datasets. To support this, we developed PC Guide (pathwaycommons.org/guide) that aims to be an online textbook for pathway analysis approaches. A current focus is pathway enrichment analysis that translate observed differences at the gene-level due to state (e.g., healthy versus diseased samples) or experimental testing (e.g., control versus treated samples) into higher-level changes at the pathway level. The *Workflows* section guides users through a step-by-step, example-driven tutorial to create Enrichment Map visualizations in Cytoscape from the analysis of RNA-seq data using Gene Set Enrichment Analysis (GSEA) (30). The *Primer* section offers intuitive descriptions of analytical techniques (e.g. Fisher’s Exact Test and GSEA) used in popular software packages and apps.

## CONCLUSION

The goal of Pathway Commons is to provide a comprehensive and user-friendly access point for researchers desiring pathway and molecular interaction information to support the analysis of biological data and the discovery of interesting relationships. Since our original report (8), the resource has expanded to include most of the widely used publicly available pathway datasets. In addition, we have increased accessibility through the development of web services, training resources and a diverse collection of end-user tools to explore and analyze the data. The PC Search web app aims to provide a unified and intelligent way to deliver relevant information and tools to users, inspired by recent additions to Google search functionality that ‘understand’ the query type to provide relevant search results (e.g. local movie times if you search for a movie name). We plan to extend the range of biological concepts recognized (e.g. drugs, metabolites, diseases) and collaborate with the community on the development of a unified and user-friendly federated search across network and pathway resources.

While PC incorporates over 20 large pathway and molecular interaction resources and over 700 of these resources are known, the vast majority of pathway resources are unfortunately no longer active or available. Further, even for the 22 databases currently integrated, much effort was required to work with data providers to create or tune BioPAX output to enable integration of the available data. For this reason, even with 700 created pathway-related databases, few additional ones will be integrated. As new databases are created, they can now use PC software components, such as Paxtools, to make available standard BioPAX formatted output. Another major barrier to pathway data access is that only a small handful of pathway and molecular interaction resources that curate data from the literature remain actively funded and they are only able to cover a relatively small part of the rapidly growing literature. To address this, the PC team is advancing text-mining technology to extract pathway information directly from the existing literature (60, 66, 67), and developing a curation support tool that empowers authors themselves to capture and share structured summaries of knowledge described in their articles. These efforts, when combined with continued expert curation, may meet the challenge of providing high-quality, computable pathway information that can be effectively searched and analyzed by the broader research community.

## DATA AVAILABILITY

All software developed as part of Pathway Commons is freely available, open-source and hosted on GitHub repositories. Software for the Pathway Commons project is hosted at github.com/PathwayCommons and software related to the BioPAX initiative is hosted at github.com/BioPAX. Users of Pathway Commons are able to provide feedback and ask questions of the development team using a discussion group (groups.google.com/forum/#!forum/pathway-commons-help). Users can submit developer feedback, file bug reports, and request new features using project-specific issue trackers (e.g., github.com/PathwayCommons/cpath2/issues or github.com/BioPAX/Paxtools/issues).

## SUPPLEMENTARY DATA

Supplementary Data are available at NAR online.

## ACKNOWLEDGEMENTS

We thank the many data providers that have collaborated with us on this project and the Google Summer of Code program for help identifying students to contribute to this project.

## REFERENCES

1. Khatri, P., Sirota, M. and Butte, A.J. (2012) Ten years of pathway analysis: current approaches and outstanding challenges. PLoS Comput. Biol., 8, e1002375.

2. Chinen, T., Kannan, A.K., Levine, A.G., Fan, X., Klein, U., Zheng, Y., Gasteiger, G., Feng, Y., Fontenot, J.D. and Rudensky, A.Y. (2016) An essential role for the IL-2 receptor in Treg cell function. Nat. Immunol., 17, 1322–1333.

3. Santos, M.A., Faryabi, R.B., Ergen, A.V., Day, A.M., Malhowski, A., Canela, A., Onozawa, M., Lee, J.-E., Callen, E., Gutierrez-Martinez, P., et al. (2014) DNA-damage-induced differentiation of leukaemic cells as an anti-cancer barrier. Nature, 514, 107–111.

4. Behan, F.M., Iorio, F., Picco, G., Gonçalves, E., Beaver, C.M., Migliardi, G., Santos, R., Rao, Y., Sassi, F., Pinnelli, M., et al. (2019) Prioritization of cancer therapeutic targets using CRISPR-Cas9 screens. Nature, 568, 511–516.

5. Sheffield, B.S., Tinker, A.V., Shen, Y., Hwang, H., Li-Chang, H.H., Pleasance, E., Ch’ng, C., Lum, A., Lorette, J., McConnell, Y.J., et al. (2015) Personalized oncogenomics: clinical experience with malignant peritoneal mesothelioma using whole genome sequencing. PloS One, 10, e0119689.

6. Mack, S.C., Witt, H., Piro, R.M., Gu, L., Zuyderduyn, S., Stütz, A.M., Wang, X., Gallo, M., Garzia, L., Zayne, K., et al. (2014) Epigenomic alterations define lethal CIMP-positive ependymomas of infancy. Nature, 506, 445–450.

7. Bader, G.D., Cary, M.P. and Sander, C. (2006) Pathguide: a pathway resource list. Nucleic Acids Res., 34, D504–506.

8. Cerami, E.G., Gross, B.E., Demir, E., Rodchenkov, I., Babur, O., Anwar, N., Schultz, N., Bader, G.D. and Sander, C. (2011) Pathway Commons, a web resource for biological pathway data. Nucleic Acids Res., 39, D685–690.

9. Demir, E., Cary, M.P., Paley, S., Fukuda, K., Lemer, C., Vastrik, I., Wu, G., D’Eustachio, P., Schaefer, C., Luciano, J., et al. (2010) The BioPAX community standard for pathway data sharing. Nat. Biotechnol., 28, 935–942.

10. Kerrien, S., Orchard, S., Montecchi-Palazzi, L., Aranda, B., Quinn, A.F., Vinod, N., Bader, G.D., Xenarios, I., Wojcik, J., Sherman, D., et al. (2007) Broadening the horizon--level 2.5 of the HUPO-PSI format for molecular interactions. BMC Biol., 5, 44.

11. Azzam, S., Schlatzer, D., Maxwell, S., Li, X., Bazdar, D., Chen, Y., Asaad, R., Barnholtz-Sloan, J., Chance, M.R. and Sieg, S.F. (2016) Proteome and Protein Network Analyses of Memory T Cells Find Altered Translation and Cell Stress Signaling in Treated Human Immunodeficiency Virus Patients Exhibiting Poor CD4 Recovery. Open Forum Infect. Dis., 3, ofw037.

12. Campbell, J., Ryan, C.J., Brough, R., Bajrami, I., Pemberton, H.N., Chong, I.Y., Costa-Cabral, S., Frankum, J., Gulati, A., Holme, H., et al. (2016) Large-Scale Profiling of Kinase Dependencies in Cancer Cell Lines. Cell Rep., 14, 2490–2501.

13. Cheng, Y., Wang, Z.-M., Tan, W., Wang, X., Li, Y., Bai, B., Li, Y., Zhang, S.-F., Yan, H.-L., Chen, Z.-L., et al. (2018) Partial loss of psychiatric risk gene Mir137 in mice causes repetitive behavior and impairs sociability and learning via increased Pde10a. Nat. Neurosci., 21, 1689–1703.

14. Grimes, M., Hall, B., Foltz, L., Levy, T., Rikova, K., Gaiser, J., Cook, W., Smirnova, E., Wheeler, T., Clark, N.R., et al. (2018) Integration of protein phosphorylation, acetylation, and methylation data sets to outline lung cancer signaling networks. Sci. Signal., 11.

15. Jia, P., Chen, X., Xie, W., Kendler, K.S. and Zhao, Z. (2018) Mega-analysis of Odds Ratio: A Convergent Method for a Deep Understanding of the Genetic Evidence in Schizophrenia. Schizophr. Bull., 10.1093/schbul/sby085.

16. Kim, S.S., Dai, C., Hormozdiari, F., van de Geijn, B., Gazal, S., Park, Y., O’Connor, L., Amariuta, T., Loh, P.-R., Finucane, H., et al. (2019) Genes with High Network Connectivity Are Enriched for Disease Heritability. Am. J. Hum. Genet., 104, 896–913.

17. Lee, S., Zhang, C., Kilicarslan, M., Piening, B.D., Bjornson, E., Hallström, B.M., Groen, A.K., Ferrannini, E., Laakso, M., Snyder, M., et al. (2016) Integrated Network Analysis Reveals an Association between Plasma Mannose Levels and Insulin Resistance. Cell Metab., 24, 172–184.

18. Müller, S., Liu, S.J., Di Lullo, E., Malatesta, M., Pollen, A.A., Nowakowski, T.J., Kohanbash, G., Aghi, M., Kriegstein, A.R., Lim, D.A., et al. (2016) Single-cell sequencing maps gene expression to mutational phylogenies in PDGF- and EGF-driven gliomas. Mol. Syst. Biol., 12, 889.

19. de Matos, P., Alcántara, R., Dekker, A., Ennis, M., Hastings, J., Haug, K., Spiteri, I., Turner, S. and Steinbeck, C. (2010) Chemical Entities of Biological Interest: an update. Nucleic Acids Res., 38, D249–254.

20. Kanehisa, M. and Goto, S. (2000) KEGG: kyoto encyclopedia of genes and genomes. Nucleic Acids Res., 28, 27–30.

21. Wishart, D.S., Feunang, Y.D., Guo, A.C., Lo, E.J., Marcu, A., Grant, J.R., Sajed, T., Johnson, D., Li, C., Sayeeda, Z., et al. (2018) DrugBank 5.0: a major update to the DrugBank database for 2018. Nucleic Acids Res., 46, D1074–D1082.

22. Wishart, D.S., Feunang, Y.D., Marcu, A., Guo, A.C., Liang, K., Vázquez-Fresno, R., Sajed, T., Johnson, D., Li, C., Karu, N., et al. (2018) HMDB 4.0: the human metabolome database for 2018. Nucleic Acids Res., 46, D608–D617.

23. Cerami, E.G., Bader, G.D., Gross, B.E. and Sander, C. (2006) cPath: open source software for collecting, storing, and querying biological pathways. BMC Bioinformatics, 7, 497.

24. Demir, E., Babur, O., Rodchenkov, I., Aksoy, B.A., Fukuda, K.I., Gross, B., Sümer, O.S., Bader, G.D. and Sander, C. (2013) Using biological pathway data with paxtools. PLoS Comput. Biol., 9, e1003194.

25. Babur, Ö., Aksoy, B.A., Rodchenkov, I., Sümer, S.O., Sander, C. and Demir, E. (2014) Pattern search in BioPAX models. Bioinforma. Oxf. Engl., 30, 139–140.

26. Dogrusoz, U., Cetintas, A., Demir, E. and Babur, O. (2009) Algorithms for effective querying of compound graph-based pathway databases. BMC Bioinformatics, 10, 376.

27. Rodchenkov, I., Demir, E., Sander, C. and Bader, G.D. (2013) The BioPAX Validator. Bioinforma. Oxf. Engl., 29, 2659–2660.

28. Le Novère, N., Hucka, M., Mi, H., Moodie, S., Schreiber, F., Sorokin, A., Demir, E., Wegner, K., Aladjem, M.I., Wimalaratne, S.M., et al. (2009) The Systems Biology Graphical Notation. Nat. Biotechnol., 27, 735–741.

29. van Iersel, M.P., Villéger, A.C., Czauderna, T., Boyd, S.E., Bergmann, F.T., Luna, A., Demir, E., Sorokin, A., Dogrusoz, U., Matsuoka, Y., et al. (2012) Software support for SBGN maps: SBGN-ML and LibSBGN. Bioinforma. Oxf. Engl., 28, 2016–2021.

30. Reimand, J., Isserlin, R., Voisin, V., Kucera, M., Tannus-Lopes, C., Rostamianfar, A., Wadi, L., Meyer, M., Wong, J., Xu, C., et al. (2019) Pathway enrichment analysis and visualization of omics data using g:Profiler, GSEA, Cytoscape and EnrichmentMap. Nat. Protoc., 14, 482–517.

31. Subramanian, A., Tamayo, P., Mootha, V.K., Mukherjee, S., Ebert, B.L., Gillette, M.A., Paulovich, A., Pomeroy, S.L., Golub, T.R., Lander, E.S., et al. (2005) Gene set enrichment analysis: a knowledge-based approach for interpreting genome-wide expression profiles. Proc. Natl. Acad. Sci. U. S. A., 102, 15545–15550.

32. Braschi, B., Denny, P., Gray, K., Jones, T., Seal, R., Tweedie, S., Yates, B. and Bruford, E. (2019) Genenames.org: the HGNC and VGNC resources in 2019. Nucleic Acids Res., 47, D786–D792.

33. UniProt Consortium (2019) UniProt: a worldwide hub of protein knowledge. Nucleic Acids Res., 47, D506–D515.

34. Luna, A., Babur, Ö., Aksoy, B.A., Demir, E. and Sander, C. (2016) PaxtoolsR: pathway analysis in R using Pathway Commons. Bioinforma. Oxf. Engl., 32, 1262–1264.

35. Gyori, B.M., Bachman, J.A., Subramanian, K., Muhlich, J.L., Galescu, L. and Sorger, P.K. (2017) From word models to executable models of signaling networks using automated assembly. Mol. Syst. Biol., 13, 954.

36. Babur, O., Dogrusoz, U., Demir, E. and Sander, C. (2010) ChiBE: interactive visualization and manipulation of BioPAX pathway models. Bioinforma. Oxf. Engl., 26, 429–431.

37. Shannon, P., Markiel, A., Ozier, O., Baliga, N.S., Wang, J.T., Ramage, D., Amin, N., Schwikowski, B. and Ideker, T. (2003) Cytoscape: a software environment for integrated models of biomolecular interaction networks. Genome Res., 13, 2498–2504.

38. Funahashi, A., Matsuoka, Y., Jouraku, A., Morohashi, M., Kikuchi, N. and Kitano, H. (2008) CellDesigner 3.5: A Versatile Modeling Tool for Biochemical Networks. Proc. IEEE, 96, 1254–1265.

39. Mi, H., Muruganujan, A., Demir, E., Matsuoka, Y., Funahashi, A., Kitano, H. and Thomas, P.D. (2011) BioPAX support in CellDesigner. Bioinforma. Oxf. Engl., 27, 3437–3438.

40. Bahceci, I., Dogrusoz, U., La, K.C., Babur, Ö., Gao, J. and Schultz, N. (2017) PathwayMapper: a collaborative visual web editor for cancer pathways and genomic data. Bioinforma. Oxf. Engl., 33, 2238–2240.

41. Franz, M., Lopes, C.T., Huck, G., Dong, Y., Sumer, O. and Bader, G.D. (2016) Cytoscape.js: a graph theory library for visualisation and analysis. Bioinforma. Oxf. Engl., 32, 309–311.

42. Benis, N., Schokker, D., Kramer, F., Smits, M.A. and Suarez-Diez, M. (2016) Building pathway graphs from BioPAX data in R. F1000Research, 5, 2414.

43. Blinov, M.L., Schaff, J.C., Vasilescu, D., Moraru, I.I., Bloom, J.E. and Loew, L.M. (2017) Compartmental and Spatial Rule-Based Modeling with Virtual Cell. Biophys. J., 113, 1365–1372.

44. Cokelaer, T., Pultz, D., Harder, L.M., Serra-Musach, J. and Saez-Rodriguez, J. (2013) BioServices: a common Python package to access biological Web Services programmatically. Bioinforma. Oxf. Engl., 29, 3241–3242.

45. Emig, D., Salomonis, N., Baumbach, J., Lengauer, T., Conklin, B.R. and Albrecht, M. (2010) AltAnalyze and DomainGraph: analyzing and visualizing exon expression data. Nucleic Acids Res., 38, W755–762.

46. Gao, J., Aksoy, B.A., Dogrusoz, U., Dresdner, G., Gross, B., Sumer, S.O., Sun, Y., Jacobsen, A., Sinha, R., Larsson, E., et al. (2013) Integrative analysis of complex cancer genomics and clinical profiles using the cBioPortal. Sci. Signal., 6, pl1.

47. Hill, S.M., Heiser, L.M., Cokelaer, T., Unger, M., Nesser, N.K., Carlin, D.E., Zhang, Y., Sokolov, A., Paull, E.O., Wong, C.K., et al. (2016) Inferring causal molecular networks: empirical assessment through a community-based effort. Nat. Methods, 13, 310–318.

48. Himmelstein, D.S., Lizee, A., Hessler, C., Brueggeman, L., Chen, S.L., Hadley, D., Green, A., Khankhanian, P. and Baranzini, S.E. (2017) Systematic integration of biomedical knowledge prioritizes drugs for repurposing. eLife, 6.

49. Huang, J.K., Carlin, D.E., Yu, M.K., Zhang, W., Kreisberg, J.F., Tamayo, P. and Ideker, T. (2018) Systematic Evaluation of Molecular Networks for Discovery of Disease Genes. Cell Syst., 6, 484–495.e5.

50. Nguyen, D.-T., Mathias, S., Bologa, C., Brunak, S., Fernandez, N., Gaulton, A., Hersey, A., Holmes, J., Jensen, L.J., Karlsson, A., et al. (2017) Pharos: Collating protein information to shed light on the druggable genome. Nucleic Acids Res., 45, D995–D1002.

51. Rouillard, A.D., Gundersen, G.W., Fernandez, N.F., Wang, Z., Monteiro, C.D., McDermott, M.G. and Ma’ayan, A. (2016) The harmonizome: a collection of processed datasets gathered to serve and mine knowledge about genes and proteins. Database J. Biol. Databases Curation, 2016.

52. Sinha, S., Song, J., Weinshilboum, R., Jongeneel, V. and Han, J. (2015) KnowEnG: a knowledge engine for genomics. J. Am. Med. Inform. Assoc. JAMIA, 22, 1115–1119.

53. Cerami, E., Demir, E., Schultz, N., Taylor, B.S. and Sander, C. (2010) Automated network analysis identifies core pathways in glioblastoma. PloS One, 5, e8918.

54. Babur, Ö., Ngo, A.T.P., Rigg, R.A., Pang, J., Rub, Z.T., Buchanan, A.E., Mitrugno, A., David, L.L., McCarty, O.J.T., Demir, E., et al. (2018) Platelet procoagulant phenotype is modulated by a p38-MK2 axis that regulates RTN4/Nogo proximal to the endoplasmic reticulum: utility of pathway analysis. Am. J. Physiol. Cell Physiol., 314, C603–C615.

55. Babur, Ö., Luna, A., Korkut, A., Durupinar, F., Siper, M.C., Dogrusoz, U., Aslan, J.E., Sander, C. and Demir, E. (2018) Causal interactions from proteomic profiles: molecular data meets pathway knowledge Systems Biology.

56. Babur, Ö., Gönen, M., Aksoy, B.A., Schultz, N., Ciriello, G., Sander, C. and Demir, E. (2015) Systematic identification of cancer driving signaling pathways based on mutual exclusivity of genomic alterations. Genome Biol., 16, 45.

57. Manning, H., O’Roak, B.J. and Babur, Ö. (2019) Mutually exclusive autism mutations point to the circadian clock and PI3K signaling pathways Genetics.

58. Korkut, A., Wang, W., Demir, E., Aksoy, B.A., Jing, X., Molinelli, E.J., Babur, Ö., Bemis, D.L., Onur Sumer, S., Solit, D.B., et al. (2015) Perturbation biology nominates upstream-downstream drug combinations in RAF inhibitor resistant melanoma cells. eLife, 4.

59. Cohen, P.R. (2015) DARPA’s Big Mechanism program. Phys. Biol., 12, 045008.

60. Valenzuela-Escárcega, M.A., Babur, Ö., Hahn-Powell, G., Bell, D., Hicks, T., Noriega-Atala, E., Wang, X., Surdeanu, M., Demir, E. and Morrison, C.T. (2018) Large-scale automated machine reading discovers new cancer-driving mechanisms. Database J. Biol. Databases Curation, 2018.

61. Raudvere, U., Kolberg, L., Kuzmin, I., Arak, T., Adler, P., Peterson, H. and Vilo, J. (2019) g:Profiler: a web server for functional enrichment analysis and conversions of gene lists (2019 update). Nucleic Acids Res., 47, W191–W198.

62. Fabregat, A., Jupe, S., Matthews, L., Sidiropoulos, K., Gillespie, M., Garapati, P., Haw, R., Jassal, B., Korninger, F., May, B., et al. (2018) The Reactome Pathway Knowledgebase. Nucleic Acids Res., 46, D649–D655.

63. Gene Ontology Consortium (2015) Gene Ontology Consortium: going forward. Nucleic Acids Res., 43, D1049–1056.

64. Merico, D., Isserlin, R., Stueker, O., Emili, A. and Bader, G.D. (2010) Enrichment map: a network-based method for gene-set enrichment visualization and interpretation. PloS One, 5, e13984.

65. Cerami, E., Gao, J., Dogrusoz, U., Gross, B.E., Sumer, S.O., Aksoy, B.A., Jacobsen, A., Byrne, C.J., Heuer, M.L., Larsson, E., et al. (2012) The cBio cancer genomics portal: an open platform for exploring multidimensional cancer genomics data. Cancer Discov., 2, 401–404.

66. Giorgi, J.M. and Bader, G.D. (2019) Towards reliable named entity recognition in the biomedical domain. Bioinforma. Oxf. Engl., 10.1093/bioinformatics/btz504.

67. Giorgi, J.M. and Bader, G.D. (2018) Transfer learning for biomedical named entity recognition with neural networks. Bioinforma. Oxf. Engl., 34, 4087–4094.

68. Bader, G.D., Betel, D. and Hogue, C.W.V. (2003) BIND: the Biomolecular Interaction Network Database. Nucleic Acids Res., 31, 248–250.

69. Breuer, K., Foroushani, A.K., Laird, M.R., Chen, C., Sribnaia, A., Lo, R., Winsor, G.L., Hancock, R.E.W., Brinkman, F.S.L. and Lynn, D.J. (2013) InnateDB: systems biology of innate immunity and beyond--recent updates and continuing curation. Nucleic Acids Res., 41, D1228–1233.

70. Chou, C.-H., Shrestha, S., Yang, C.-D., Chang, N.-W., Lin, Y.-L., Liao, K.-W., Huang, W.-C., Sun, T.-H., Tu, S.-J., Lee, W.-H., et al. (2018) miRTarBase update 2018: a resource for experimentally validated microRNA-target interactions. Nucleic Acids Res., 46, D296–D302.

71. Davis, A.P., Grondin, C.J., Johnson, R.J., Sciaky, D., King, B.L., McMorran, R., Wiegers, J., Wiegers, T.C. and Mattingly, C.J. (2017) The Comparative Toxicogenomics Database: update 2017. Nucleic Acids Res., 45, D972–D978.

72. Giurgiu, M., Reinhard, J., Brauner, B., Dunger-Kaltenbach, I., Fobo, G., Frishman, G., Montrone, C. and Ruepp, A. (2019) CORUM: the comprehensive resource of mammalian protein complexes-2019. Nucleic Acids Res., 47, D559–D563.

73. Hornbeck, P.V., Zhang, B., Murray, B., Kornhauser, J.M., Latham, V. and Skrzypek, E. (2015) PhosphoSitePlus, 2014: mutations, PTMs and recalibrations. Nucleic Acids Res., 43, D512–520.

74. Kandasamy, K., Mohan, S.S., Raju, R., Keerthikumar, S., Kumar, G.S.S., Venugopal, A.K., Telikicherla, D., Navarro, J.D., Mathivanan, S., Pecquet, C., et al. (2010) NetPath: a public resource of curated signal transduction pathways. Genome Biol., 11, R3.

75. Keshava Prasad, T.S., Goel, R., Kandasamy, K., Keerthikumar, S., Kumar, S., Mathivanan, S., Telikicherla, D., Raju, R., Shafreen, B., Venugopal, A., et al. (2009) Human Protein Reference Database--2009 update. Nucleic Acids Res., 37, D767–772.

76. Mi, H., Huang, X., Muruganujan, A., Tang, H., Mills, C., Kang, D. and Thomas, P.D. (2017) PANTHER version 11: expanded annotation data from Gene Ontology and Reactome pathways, and data analysis tool enhancements. Nucleic Acids Res., 45, D183–D189.

77. Orchard, S., Ammari, M., Aranda, B., Breuza, L., Briganti, L., Broackes-Carter, F., Campbell, N.H., Chavali, G., Chen, C., del-Toro, N., et al. (2014) The MIntAct project--IntAct as a common curation platform for 11 molecular interaction databases. Nucleic Acids Res., 42, D358–363.

78. Pico, A.R., Kelder, T., van Iersel, M.P., Hanspers, K., Conklin, B.R. and Evelo, C. (2008) WikiPathways: pathway editing for the people. PLoS Biol., 6, e184.

79. Romero, P., Wagg, J., Green, M.L., Kaiser, D., Krummenacker, M. and Karp, P.D. (2005) Computational prediction of human metabolic pathways from the complete human genome. Genome Biol., 6, R2.

80. Salwinski, L., Miller, C.S., Smith, A.J., Pettit, F.K., Bowie, J.U. and Eisenberg, D. (2004) The Database of Interacting Proteins: 2004 update. Nucleic Acids Res., 32, D449–451.

81. Schaefer, C.F., Anthony, K., Krupa, S., Buchoff, J., Day, M., Hannay, T. and Buetow, K.H. (2009) PID: the Pathway Interaction Database. Nucleic Acids Res., 37, D674–679.

82. Stark, C., Breitkreutz, B.-J., Reguly, T., Boucher, L., Breitkreutz, A. and Tyers, M. (2006) BioGRID: a general repository for interaction datasets. Nucleic Acids Res., 34, D535–539.

83. Thiele, I., Swainston, N., Fleming, R.M.T., Hoppe, A., Sahoo, S., Aurich, M.K., Haraldsdottir, H., Mo, M.L., Rolfsson, O., Stobbe, M.D., et al. (2013) A community-driven global reconstruction of human metabolism. Nat. Biotechnol., 31, 419–425.

84. Wrzodek, C., Büchel, F., Ruff, M., Dräger, A. and Zell, A. (2013) Precise generation of systems biology models from KEGG pathways. BMC Syst. Biol., 7, 15.

85. Yamamoto, S., Sakai, N., Nakamura, H., Fukagawa, H., Fukuda, K. and Takagi, T. (2011) INOH: ontology-based highly structured database of signal transduction pathways. Database J. Biol. Databases Curation, 2011, bar052.

