## SUPPLEMENTAL DATA for "Pathway Commons: 2019 Update"

### SUPPLEMENTARY DATA

#### Table of Contents

**Purpose of this document** 1

**Description of Simple Interaction Format (SIF)** 1

- Binary interaction patterns in a BioPAX model 1
  - controls-state-change-of 2
    - via direct control 2
    - as a participant 3
    - as a controlling input 4
    - through controller small molecule 4
    - through binding small molecule 5
    - through degradation 6
  - controls-phosphorylation-of 7
  - controls-transport-of 8
  - controls-expression-of 9
    - through TemplateReaction 9
    - through Conversion 10
  - catalysis-precedes 10
  - in-complex-with 11
  - interacts-with 12
  - neighbor-of 12
  - consumption-controlled-by 13
  - controls-production-of 14
  - controls-transport-of-chemical 14
  - chemical-affects 15
    - through binding 15
    - through a control 16
  - reacts-with 16
  - used-to-produce 17
- Example pathways using binary interactions 17
  - The Citric Acid Cycle 18
    - TCA cycle from wikipathways 18
    - Binary network from Pathway Commons 18
  - BMP signaling 19
    - Wiki pathways 20
    - Binary network from Pathway Commons 20
  - AMPK signaling 21

Wiki pathways 21
Binary network from Pathway Commons 21
ABC-mediated transport 23
Reactome 23
Binary network from Pathway Commons 23

### Purpose of this document

BioPAX can encode very detailed information about biological processes. Analysis of this data, however, can be complicated as one needs to consider a wide array of n-ary relationships, different states of entities and generic classes of molecules. We provide the data in an alternative format, called SIF, that is much simpler in structure and encodes some essential information in the BioPAX data. We search certain graph patterns in the BioPAX structure to detect SIF relations. This document describes those graph patterns and provides examples.

### Description of Simple Interaction Format (SIF)

#### Binary interaction patterns in a BioPAX model

We define the below binary interaction types for capturing certain relations in a BioPAX model:

controls-state-change-of
controls-phosphorylation-of
controls-transport-of
controls-expression-of
catalysis-precedes
in-complex-with
interacts-with
consumption-controlled-by
controls-production-of
controls-transport-of-chemical
chemical-affects
reacts-with
used-to-produce

An interaction type can be captured with more than one pattern in a BioPAX model. For instance we use 6 different patterns to represent the controls-state-change-of relation.

In the current proposal, an interaction type can have either a protein or a small molecule as source or target. For a specific interaction type, the type of the source or target is fixed. For instance the first 8 interaction types in the above list are always between proteins. The next 4 interaction types are always between a protein and a small molecule. The last 2 are always between small molecules.

Proteins are identified with HGNC gene symbols. Even though a gene can produce more than one protein, and UniProt IDs are better suited for identifying proteins, we sacrifice this resolution to create readable node names. We identify small molecules with the display names of their SmallMoleculeReference.

#### controls-state-change-of

This relation defines directed interactions between proteins. It means the first protein has an effect on an event that changes the state of the second protein. Many signaling relations that are transmitted through protein modifications are captured with this relation.

This relation has 2 sub-relations: controls-phosphorylation-of and controls-transport-of.

##### via direct control

The first protein is a controller of an interaction that modifies the second protein post-translationally. The controller protein is also not involved in the reaction as a participant.

Example: A complex of Ephrin transfers GDP to HRAS, and a complex of AR transfers GTP to HRAS.

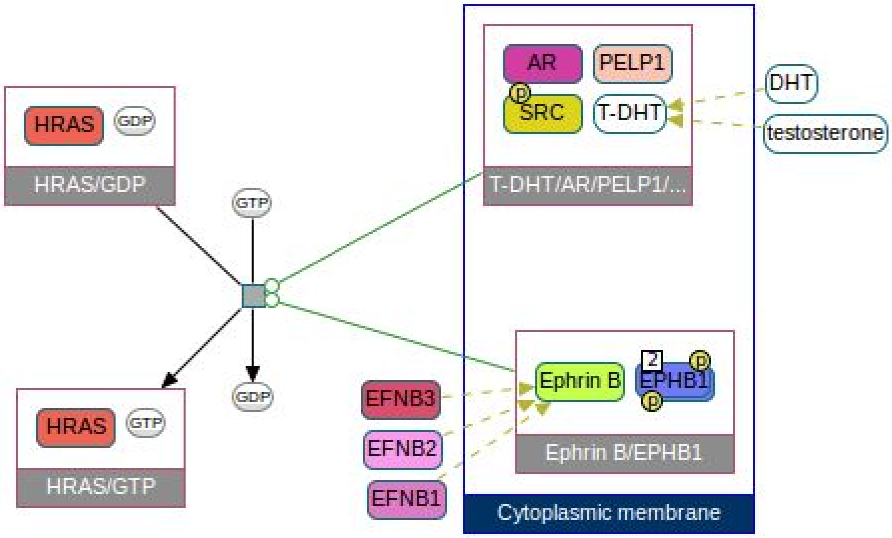


Using the data above, we generate the controls-state-change-of relations below.

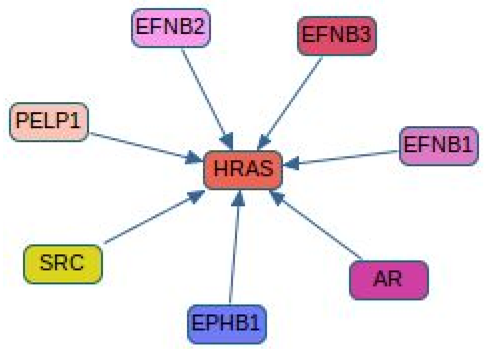


**as a participant**

Sometimes Reactome prefers an odd way of showing controllers. They make the controller both input and output using the same PhysicalEntity. This pattern captures it.

Example: An Interferon-JAK-Stat complex phosphorylates STAT1.

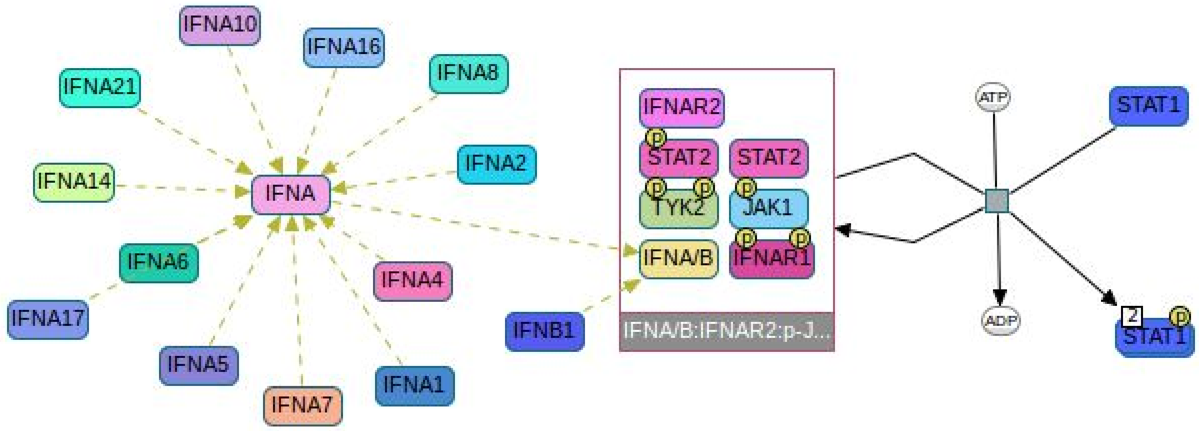


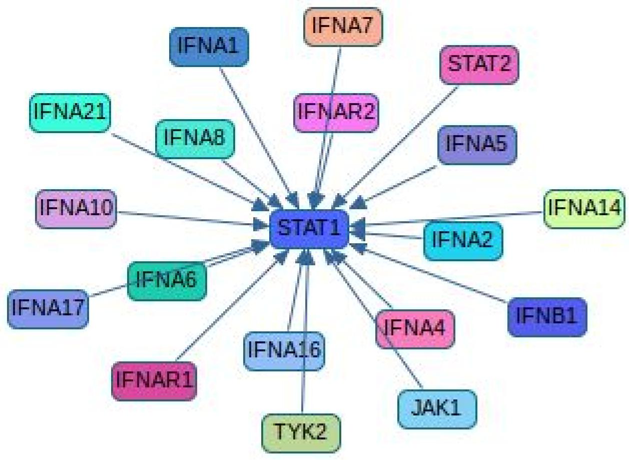

##### as a controlling input

This is another Reactome pattern for controls-state-change-of. The controller molecule is also an input to the conversion; and it is also associated with output with the same non-generic physical entity. The affected protein is represented with different simple physical entities on both sides of the reaction, where each simple physical entity is associated with only one side.

<u>Example:</u> SYK is phosphorylated by a complex containing LCK and FYN. SYK is also a member of this complex.

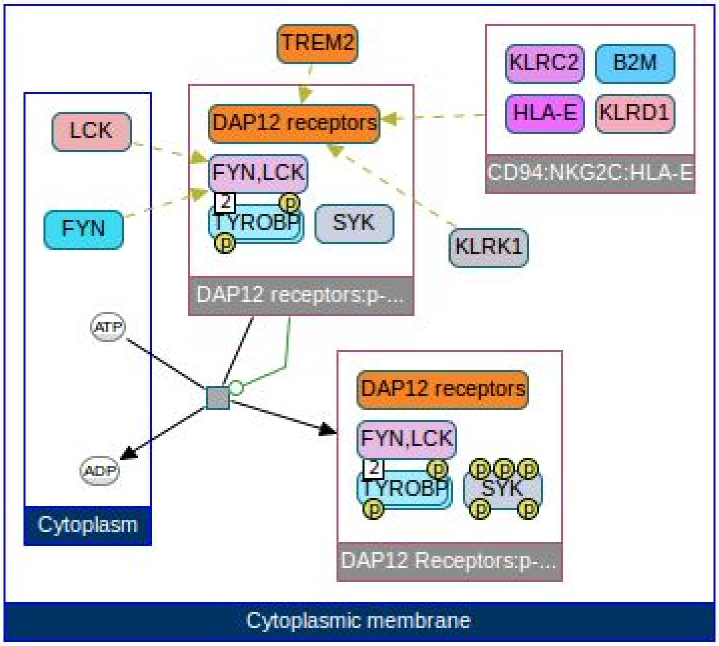


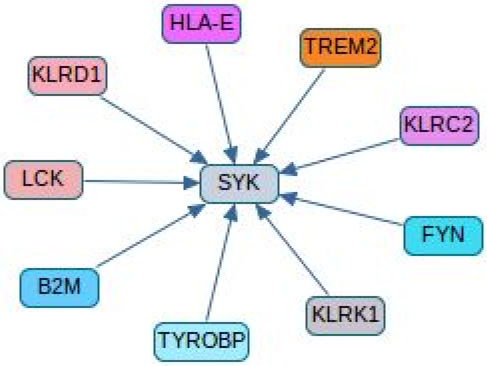


##### through controller small molecule

First protein controls a conversion that produces a small molecule. This small molecule controls a conversion that modifies the second protein. The first reaction should be changing the concentration of the linker small molecule. This means the linker small molecule should not be a blacklisted ubiquitous molecule.

Example: Activated SMPD2 catalyzes a reaction that turns sphingomyelin into ceramide. Ceramide activates another reaction that activates JNK family proteins by phosphorylation.

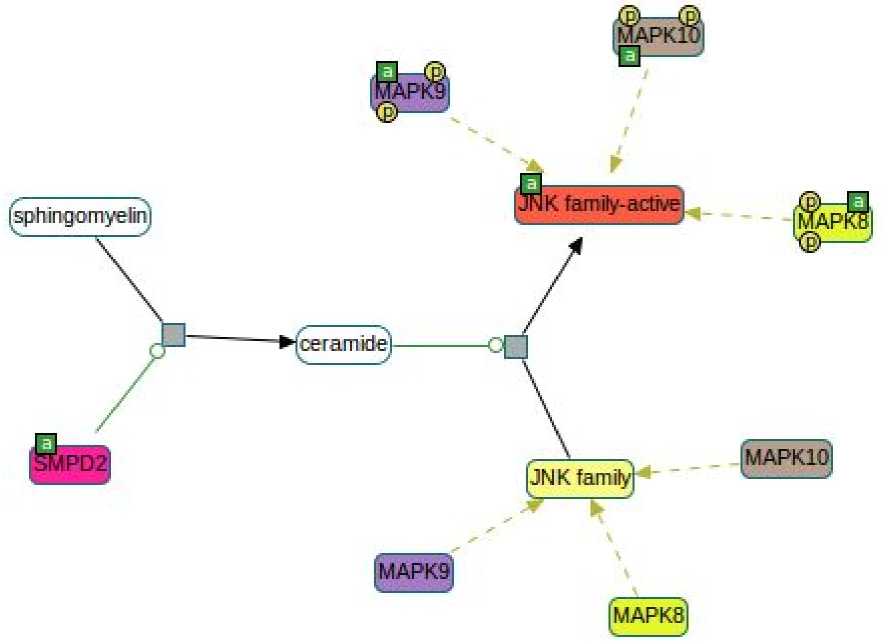


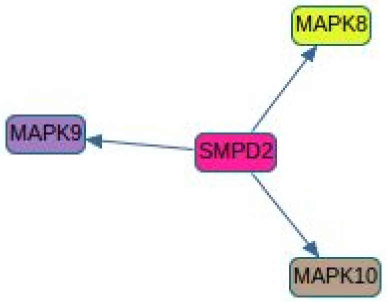


**through binding small molecule**

First protein controls a reaction that produces a small molecule. The small molecule forms a complex with the second protein. The first reaction should be changing the concentration of the linker small molecule.

Example: The PIK3CA complex catalyses production of PIP3. PIP3 binds to an AKT1 complex.

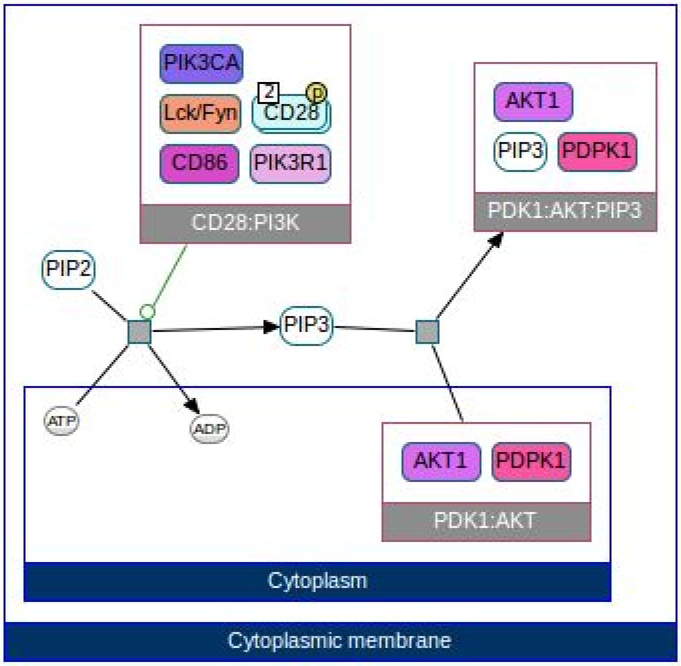


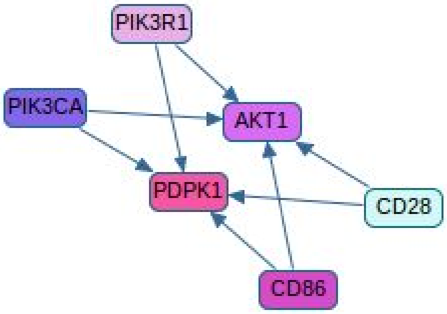


**through degradation**

First protein is controlling a reaction that degrades the second protein.

Example: Smurf proteins control degradation of SMADs.

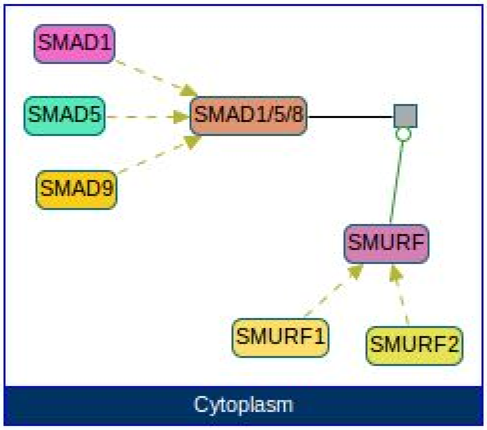


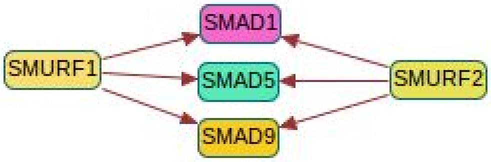


##### controls-phosphorylation-of

This is directed relation between proteins in which the first protein changes the phosphorylation status of the second protein. This is a sub-relation of controls-state-change-of. That means every controls-phosphorylation-of relation is also a controls-state-change-of relation. This relation can be useful for researchers studying phospho-proteomics.

Example: RAF phosphorylates MEK.

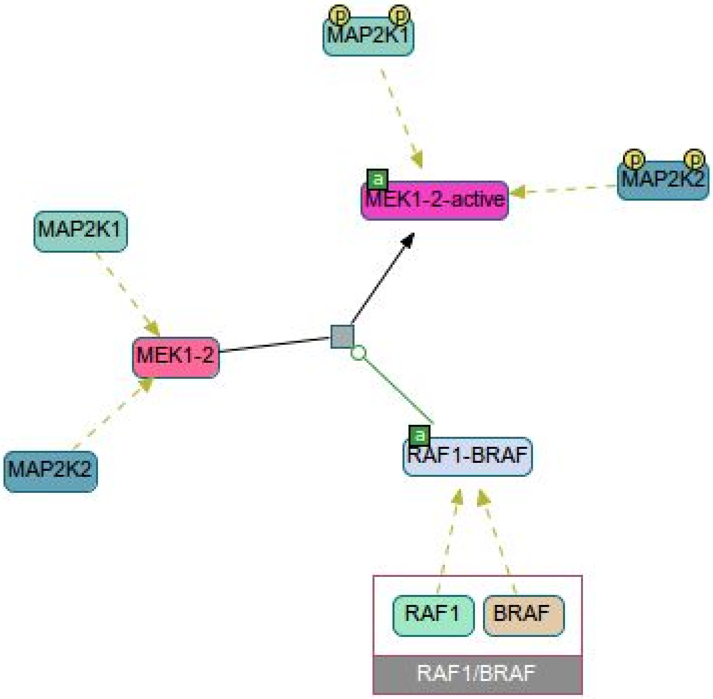


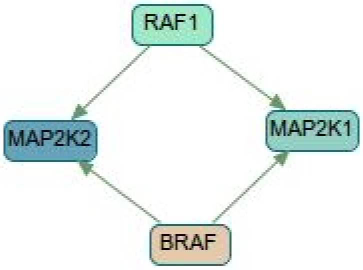


**controls-transport-of**

This is a directed relation between proteins in which the first protein controls a reaction that changes the cellular location of the second protein. Changing cellular location from null to something and from something to null does not qualify as transportation. It has to be between two non-null values. This relation type is a sub-type of controls-state-change-of. That means every controls-transport-of relation is also a controls-state-change-of relation. This relation can be useful for researchers studying cell trafficking.

<u>Example:</u> PTPN11 makes GRB2 dissociate from a membrane bound complex, hence, changes its location to cytoplasm.

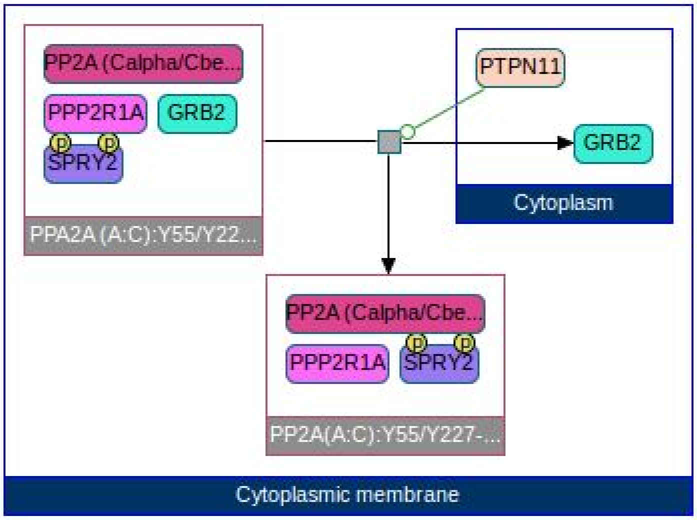


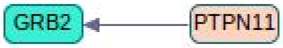


**controls-expression-of**

Defines a directed relation between two proteins where the first protein controls expression of the second protein. This relation can be useful for relating an alteration of a gene to another gene's expression change.

**through TemplateReaction**

The right way of modeling an expression in BioPAX is using a TemplateReaction. This pattern is composed of a controller, the controlled template reaction and the product.

Example: A complex of CREBBP activates expression of VEGFA.

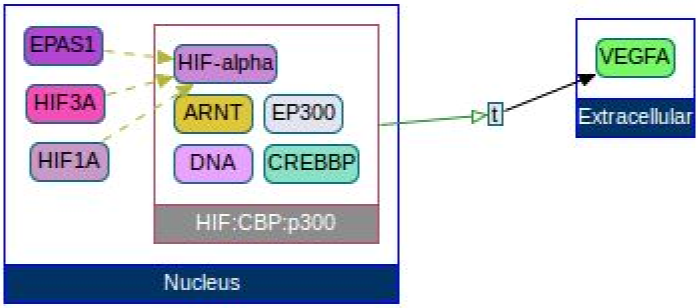

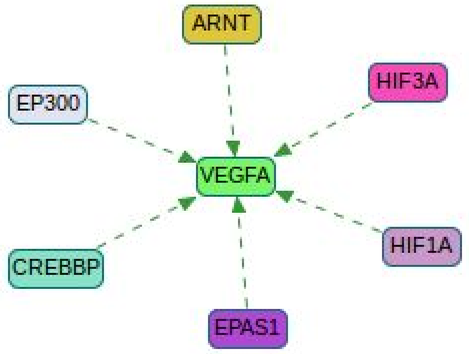


**through Conversion**

Some databases are still using a Conversion to model expression (NCI). These are conversions with only one right-participant, and no left-participant. This second pattern captures those expressions modeled in wrong way.

Example: RB1 bound TP53 inhibits TERT expression.

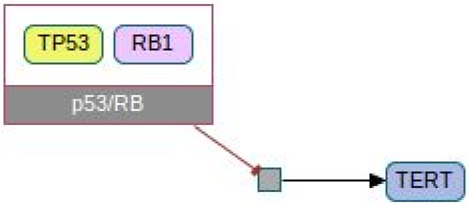


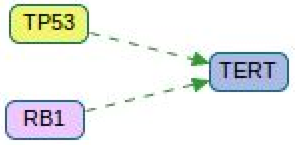


##### catalysis-precedes

Defines directed relations between two proteins that control consecutive reactions. The first protein controls a reaction whose output molecule is input to another reaction. This other reaction is controlled by the second protein. Both reactions are capable of changing concentration of the linker small molecule. The reactions cannot be identical or reverse of each other. Unshared participants at the non-facing sides of two reactions should contain at least one non-ubiquitous small molecule.

This relation can be useful for recovering metabolic cycles in the data.

Example: CYP2J2 turns arachidonate into epoxide, and epoxide hydrolyases turn the epoxide into a diol.

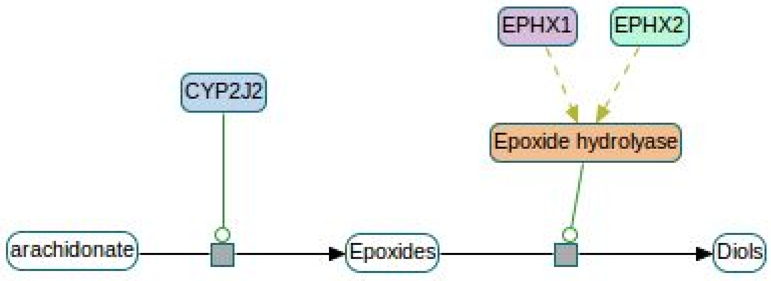


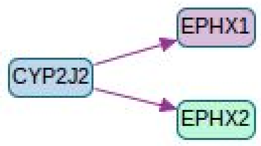


**in-complex-with**

Two proteins appear as components of the same complex. This excludes the pairs of components that cannot be members of the complex at the same time. In an ideal world, this undirected relation should reproduce the whole PPI network.

Example: WWTR1 and YAP1, separately, forms complexes with members of TEAD family. Notice that this relation is not defined between members of TEAD proteins.

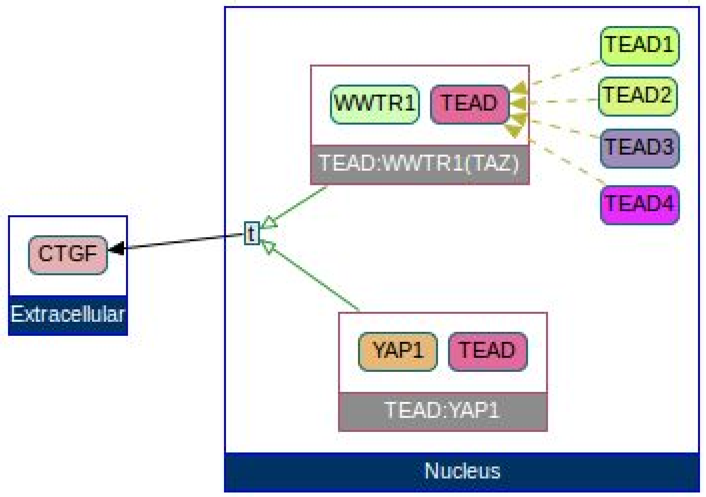

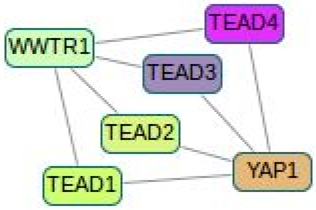


#### interacts-with

This is an undirected relation between participant proteins of a MolecularInteraction. This relation is for capturing protein interaction data (or data at the same level of simplicity), like data from HPRD.

#### neighbor-of

Two proteins are related to the same interaction (any kind), through being inputs, outputs, or controllers. This relation is useful when we need to get a list of related proteins, but don't care about the nature of the relation, and all we need is a vague neighborhood. This pattern tends to generate a lot of edges, hence, even small networks look like a hairball.

Relation to first version: This is a new interaction type. It resembles the previous INTERACTS_WITH, however this one also includes controllers of the interactions.

Example: A cyclin/CDK complex makes RB to dissociate from E2F complex. Resulting binary network is a clique with all 12 non-generic proteins involved in the process.

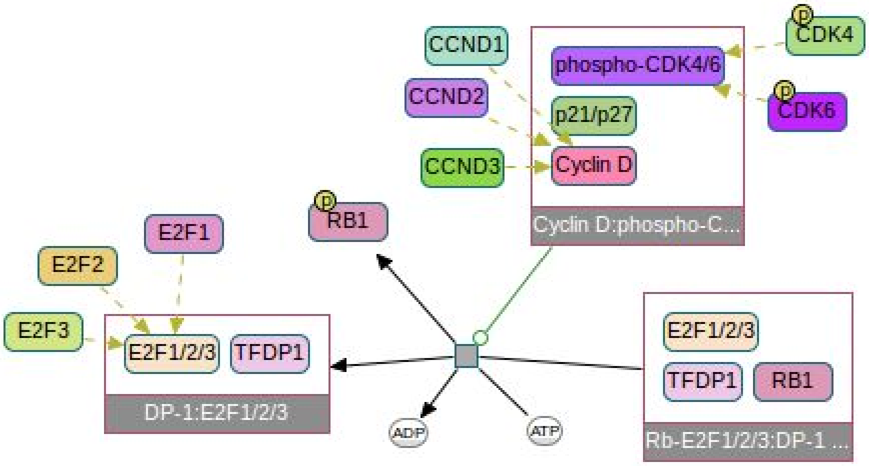

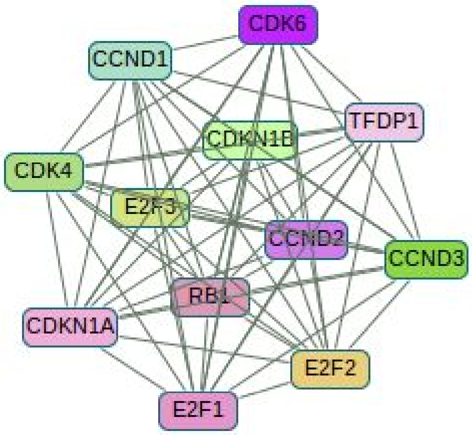


**consumption-controlled-by**

This is a directed relation from a small molecule to a protein. The protein is controlling the consumption of the small molecule. The reaction is supposed to change the concentration of the small molecule. The consumed small molecule (input) cannot also be an output. This relation can be useful for studying metabolic events.

Example: Below is part of the TCA cycle, where FH catalyses Fumaric acid and L-Malate, and MDH2 catalyses L-Malate and Oxaloacetate.

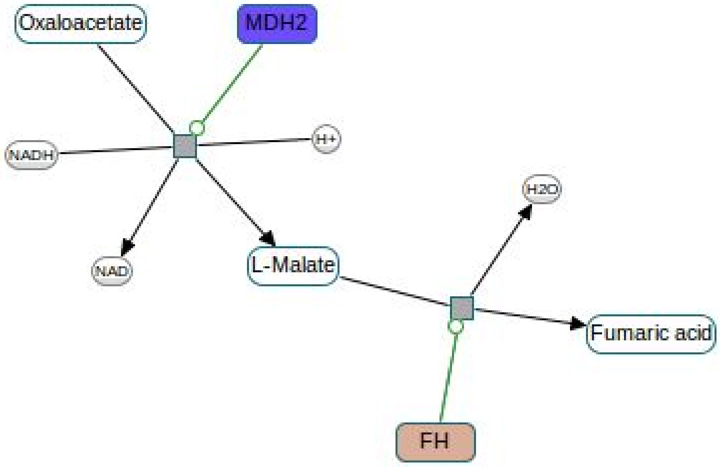

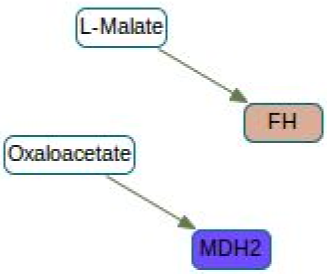


##### controls-production-of
This is a directed relation from a protein to a small molecule. The protein is controlling the production of the small molecule. The reaction is supposed to change the concentration of the small molecule. The produced small molecule (output) cannot also be an input. This relation can be useful for studying metabolic events.

Example: Consider same reactions above with MDH2 and FH. This pattern produces below interactions.

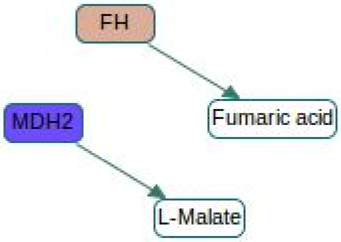


When consumption-controlled-by and controls-production-of used at the same time, we get a graph that can show the flow of the metabolic events as shown below. But users should be aware that the proteins may not be always promoting the flow, but can also be inhibiting it.

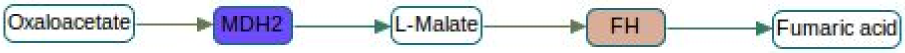


##### controls-transport-of-chemical
Directed relation from a protein to a non-generic small molecule, where the protein controls a reaction that changes the cellular location of the small molecule. This relation implies the concentration of the small molecule changes in at least one cellular location.

Example: SAR1B controls transportation of a Complex of APO proteins bound with cholesterols, triglycerides, phospholipids, and cholesterol esters.

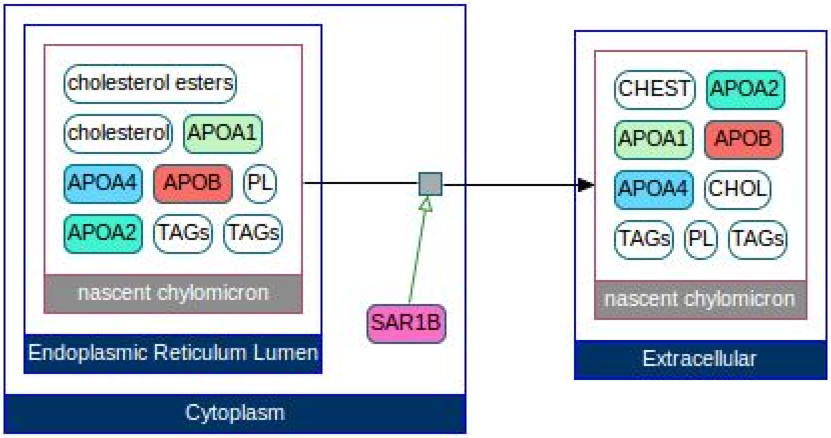


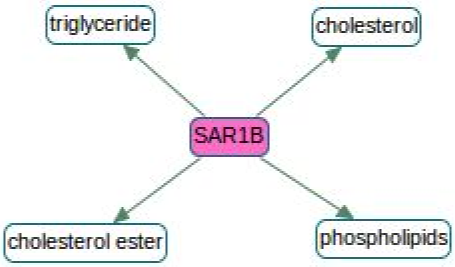


##### chemical-affects

This is a directed relation from a small molecule to a protein. The small molecule shows signs of affecting either activity or the state of the protein. This relation is essential for showing drugs in networks.

###### through binding

Similar to in-complex-with pattern, but this one is directed and it is from the small molecule to the protein.

Example: Testosterone and DHT affects activity of AR, SRC and PELP1 through binding.

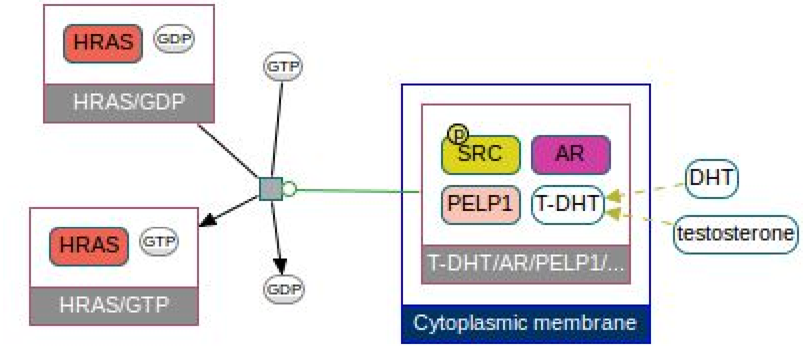


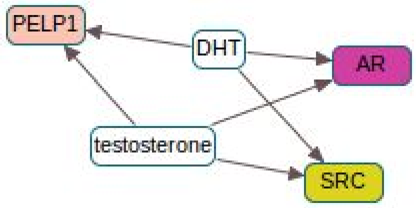


**through a control**
A small molecule is controlling an interaction and a protein is a participant of this interaction.

Example: Testosterone and DHT affects state of HRAS by controlling a reaction that modifies it (refer to the same BioPAX graph above).

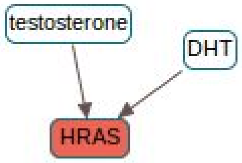


#### reacts-with

This is an undirected relation between two small molecules that are substrates to the same biochemical reaction. None of the molecules are also in products.

Example: UDPglucuronate reacts with Phenol to produce O-glucuronide and UDP.

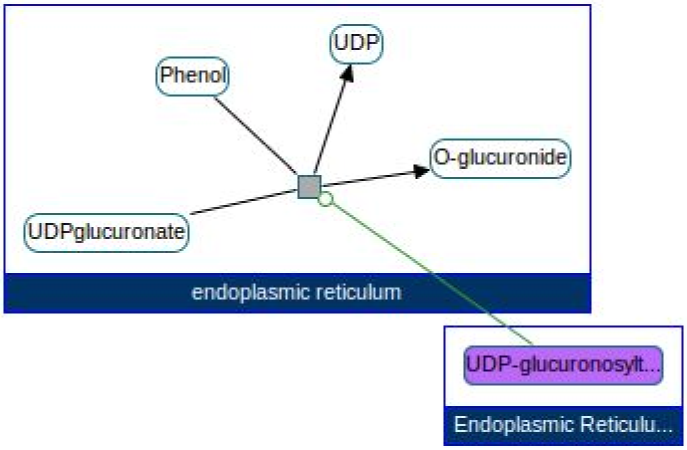


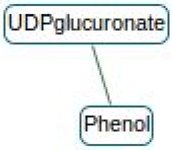


**used-to-produce**

This is a directed relation between small molecules. First one is a substrate of a biochemical reaction that produces the second molecule. Both molecules are only on one side of the reaction.

Example: Refer to the same example above where UPDglucuronate and Phenol is used to produce O-glucuronide and UDP.

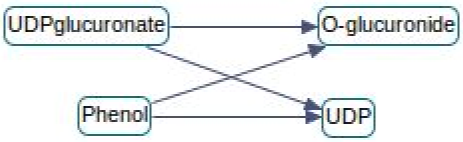


#### Example pathways using binary interactions

In this section, we try to redraw some popular pathway diagrams using binary interactions and Pathway Commons data. To produce these binary network, we execute a paths-between query in the large binary network derived from Pathway Commons, using one or two of the interaction types.

#### The Citric Acid Cycle

##### TCA cycle from wikipathways

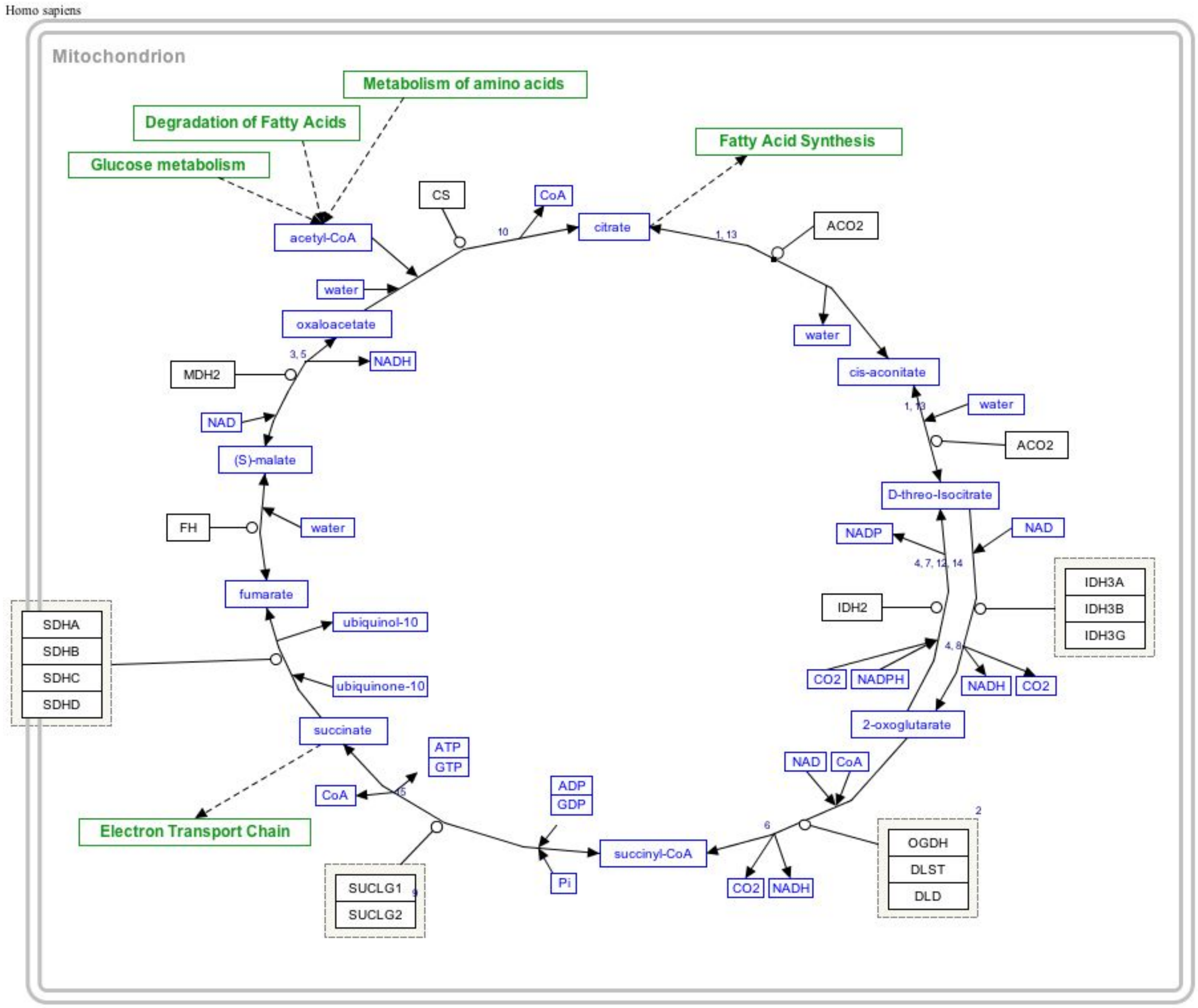


##### Binary network from Pathway Commons

With catalysis-precedes and in-complex-with interactions:

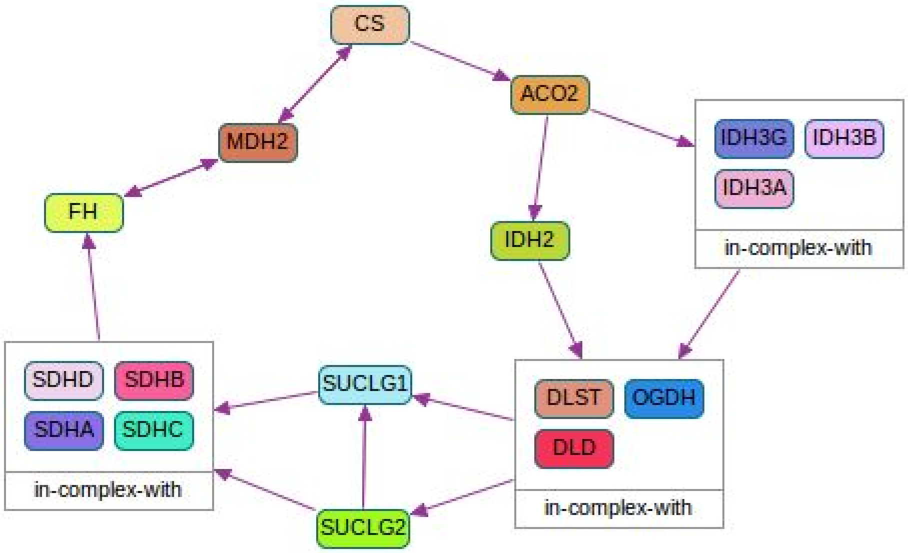


With consumption-controlled-by and controls-production-of interactions:

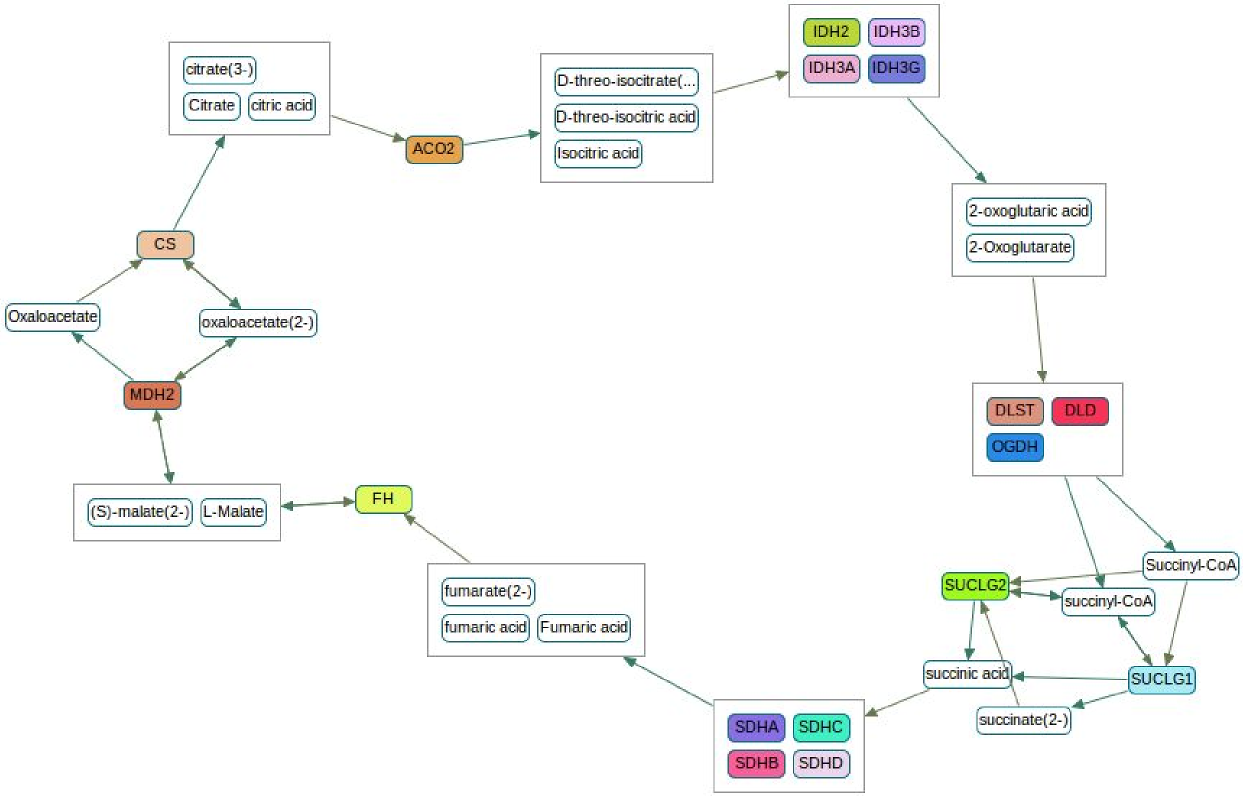


#### BMP signaling

#### Wiki pathways

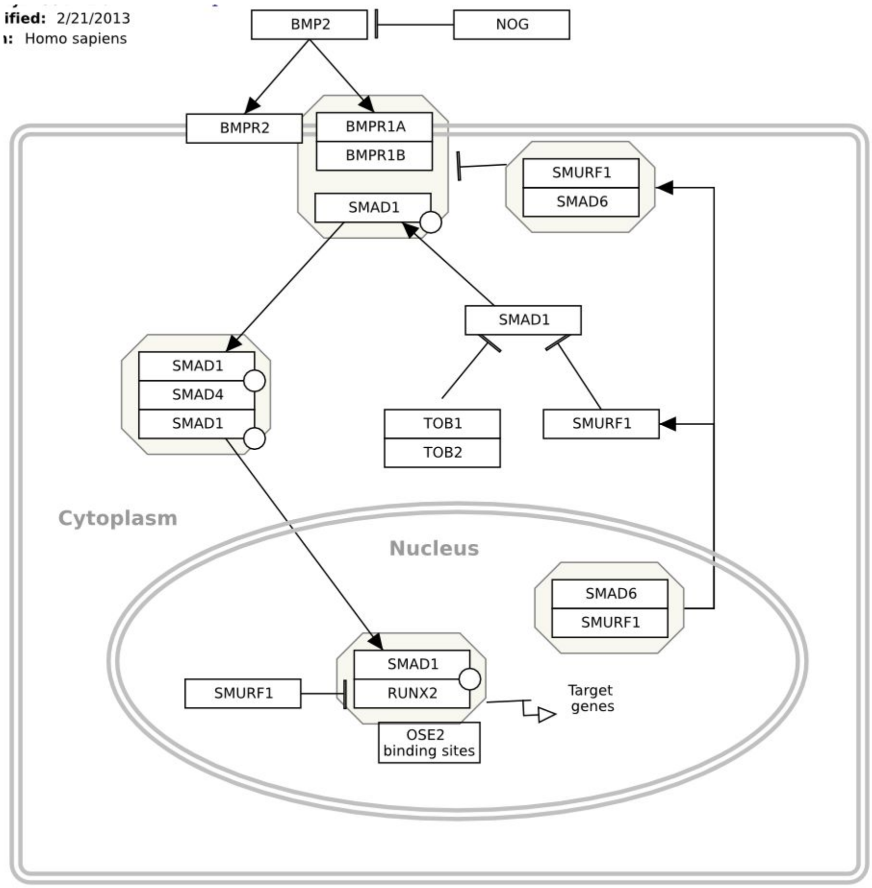


#### Binary network from Pathway Commons

With controls-state-change-of and controls-degradation-of interactions:

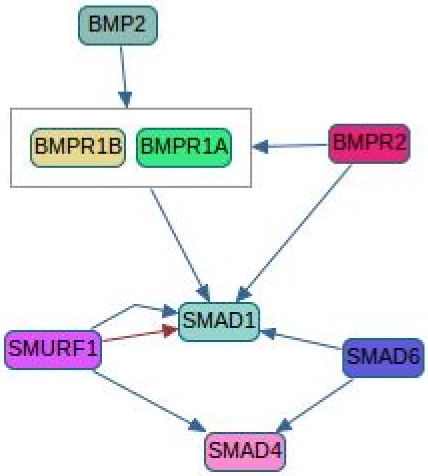

#### AMPK signaling

##### Wiki pathways

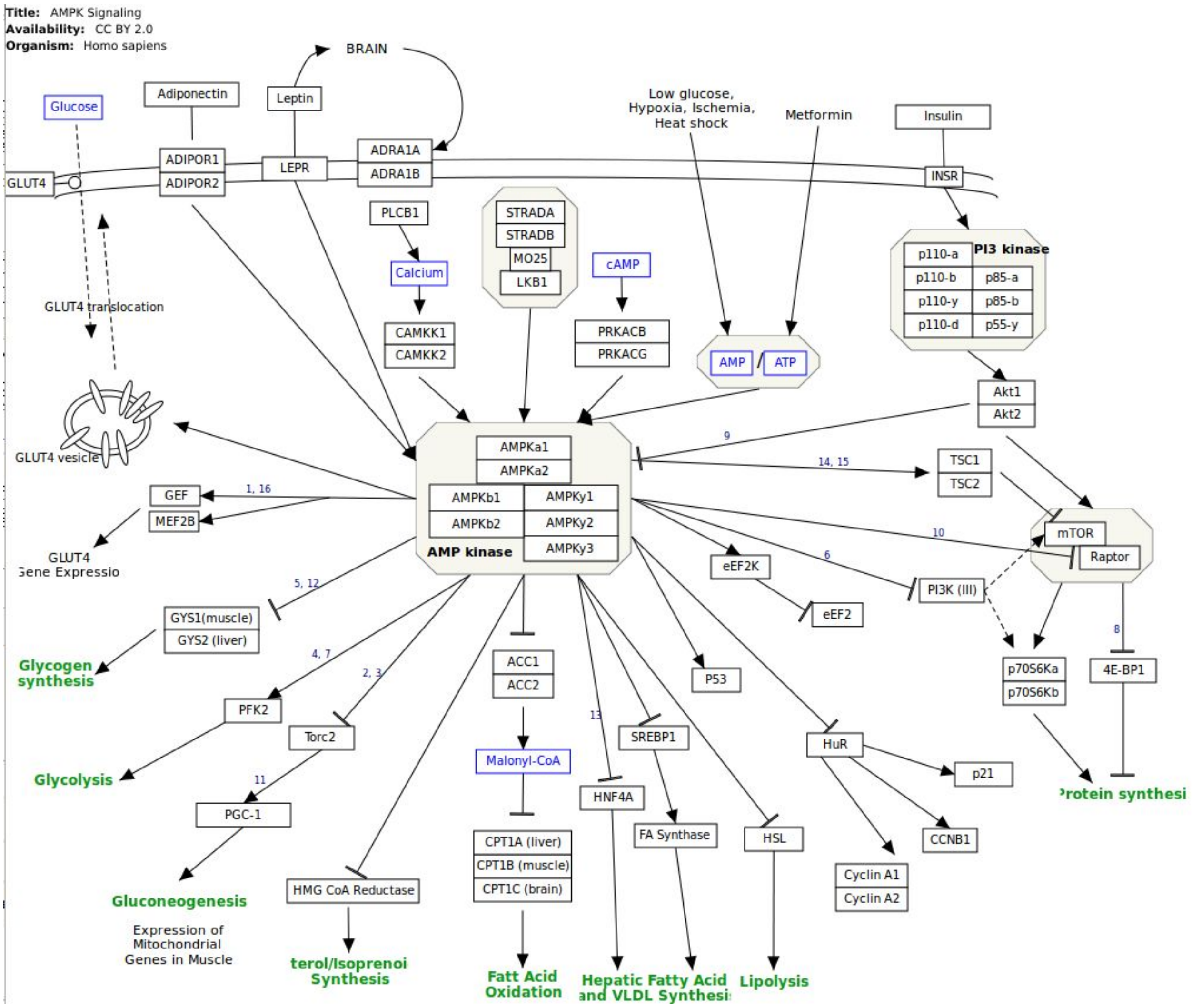


##### Binary network from Pathway Commons

With controls-state-change-of and controls-expression-of interactions (AMPK proteins are the ones starting with PRKA):

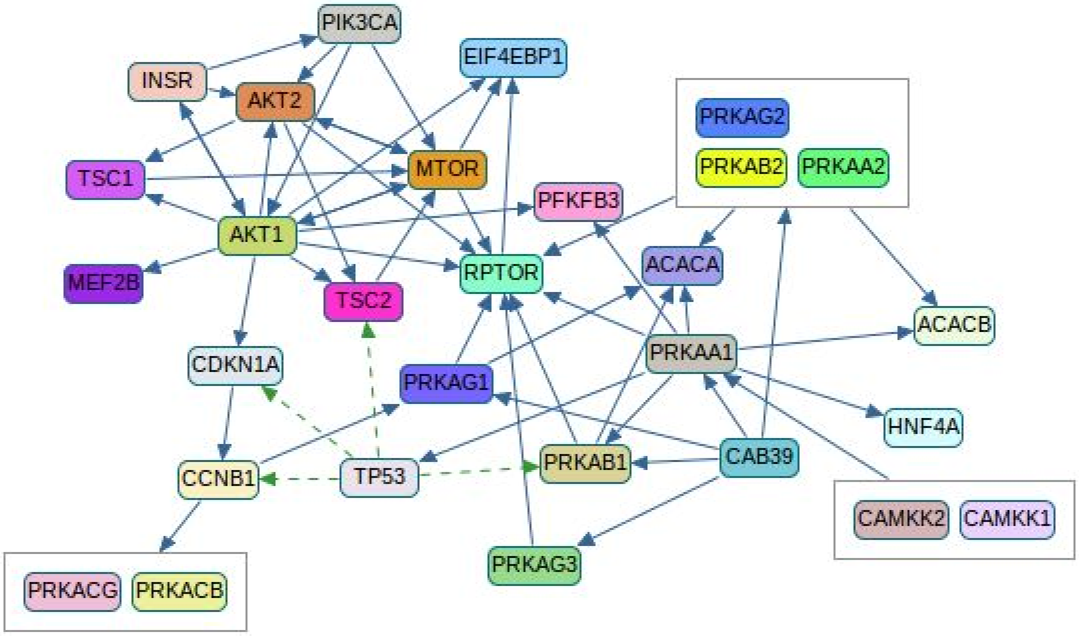


With interacts-with interactions:

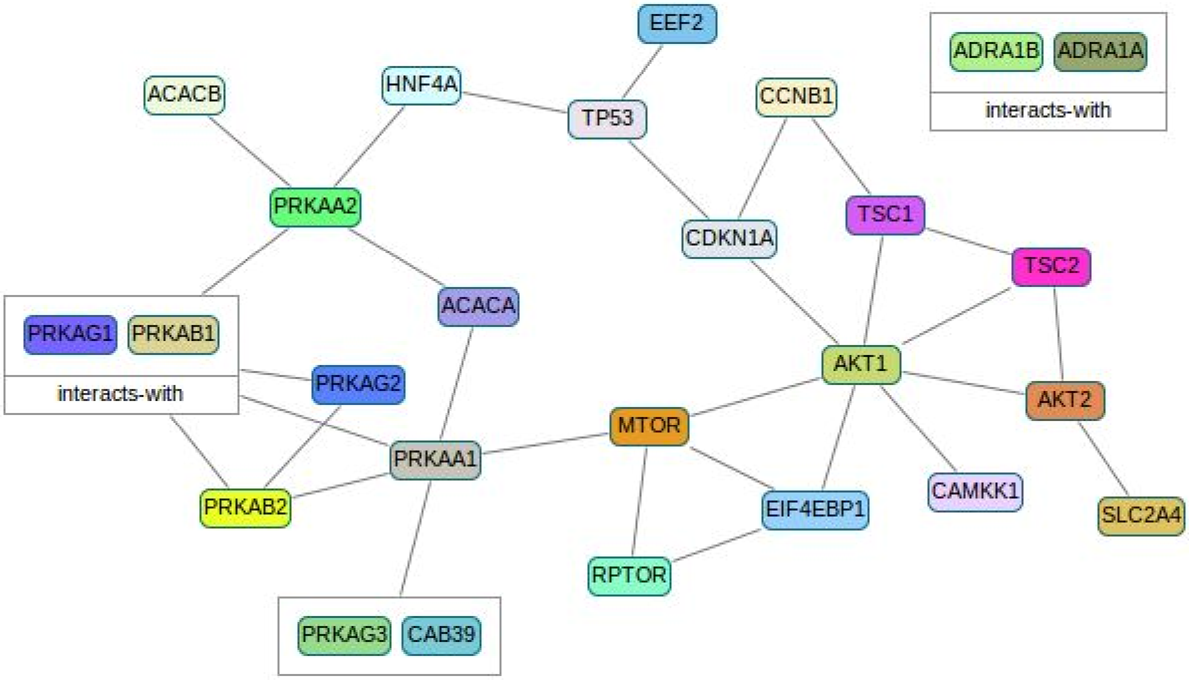

#### ABC-mediated transport

##### Reactome

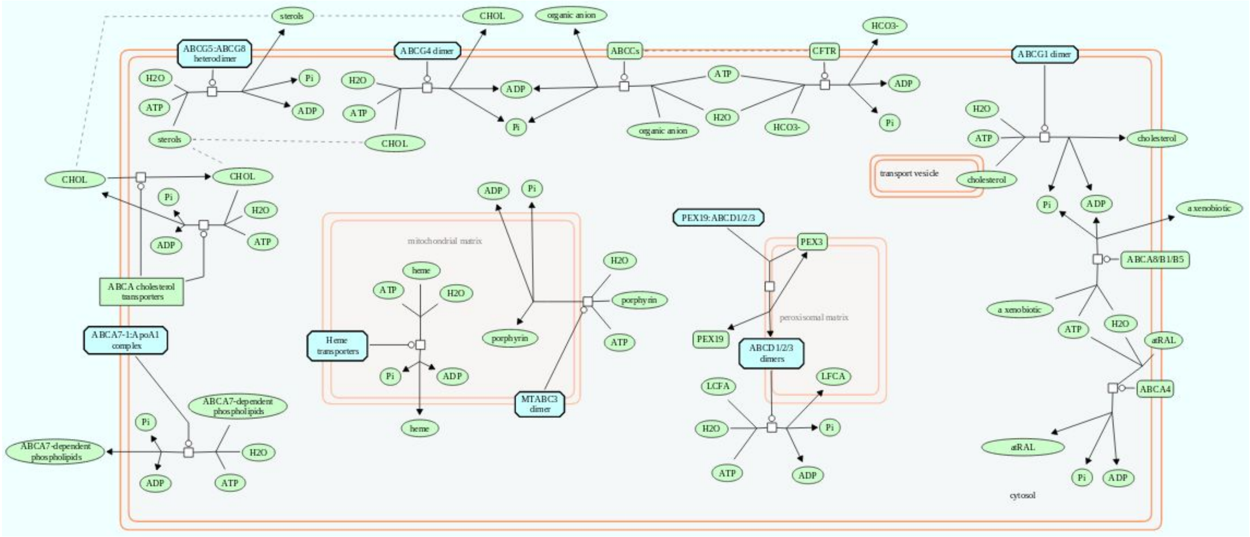


Binary network from Pathway Commons

With controls-transport-of-chemical interactions:

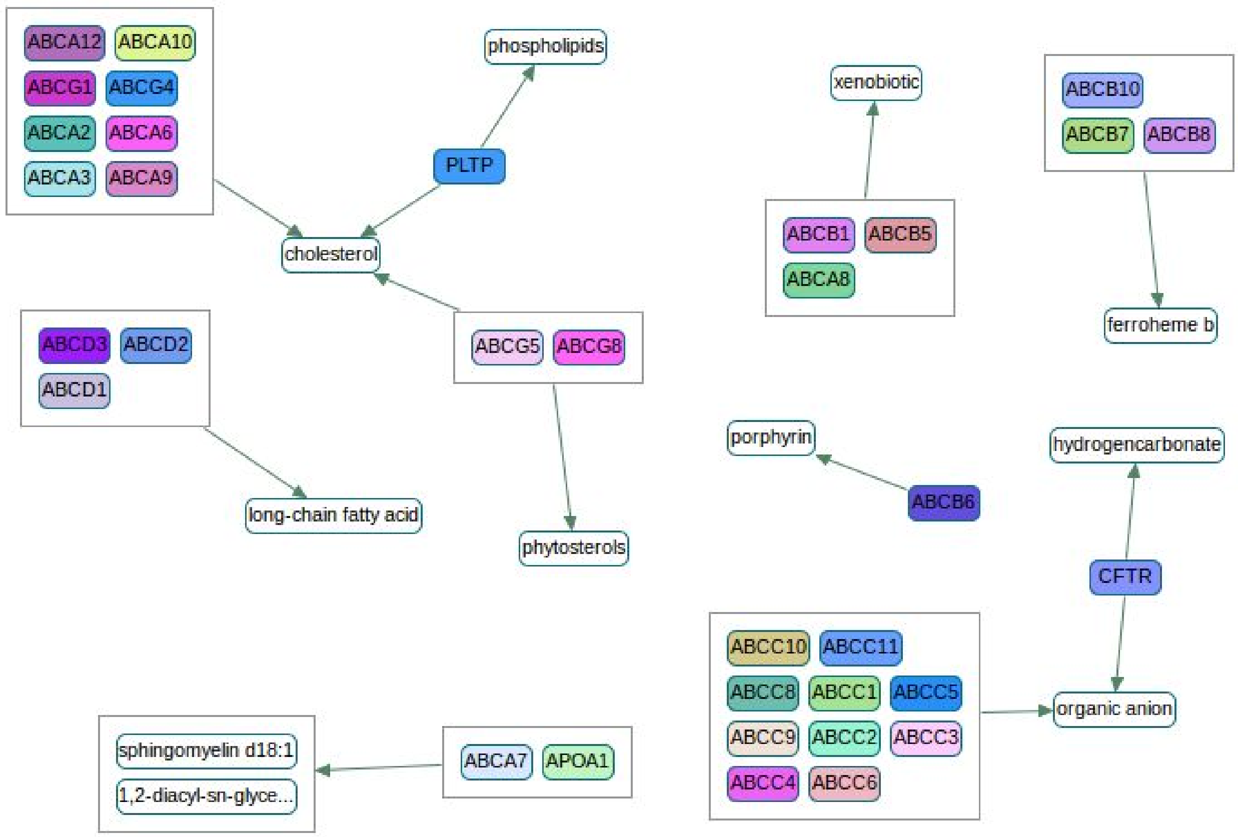
ABCA12
ABCA10
ABCG1
ABCG4
ABCA2
ABCA6
ABCA3
ABCA9
phospholipids
PLTP
cholesterol
xenobiotic
ABCB10
ABCB7
ABCB8
ABCB1
ABCB5
ABCA8
ferroheme b
ABCD3
ABCD2
ABCD1
ABCG5
ABCG8
long-chain fatty acid
phytosterols
porphyrin
ABCB6
hydrogencarbonate
CFTR
ABCC10
ABCC11
ABCC8
ABCC1
ABCC5
ABCC9
ABCC2
ABCC3
ABCC4
ABCC6
organic anion
sphingomyelin d18:1
1,2-diacyl-sn-glyce...
ABCA7
APOA1
